# Relative water content consistently predicts drought mortality risk in seedling populations with different morphology, physiology, and times to death

**DOI:** 10.1101/2020.12.08.416917

**Authors:** Gerard Sapes, Anna Sala

**Author notes:** **Corresponding Author**: Gerard Sapes, Department of Ecology, Evolution, and Behavior, University of Minnesota, 1987 Upper Buford Circle 55108, Saint Paul, Minnesota, USA.

## Abstract

Predicted increases in forest drought mortality highlight the need for predictors of incipient drought-induced mortality (DIM) risk that enable proactive large-scale management. Such predictors should be consistent across plants with varying morphology and physiology. Because of their integrative nature, indicators of water status are promising candidates for real time monitoring of DIM, particularly if they standardize morphological differences among plants. We assessed the extent to which differences in morphology and physiology between *Pinus ponderosa* populations influence time to mortality and the predictive power of key indicators of DIM risk. Time to incipient mortality differed between populations but occurred at the same relative water content (RWC) and water potential (WP). RWC and WP were accurate predictors of drought mortality risk. These results highlight that variables related to water status capture critical thresholds during DIM and the associated dehydration processes. Both WP and RWC are promising candidates for large-scale assessments of DIM risk. RWC is of special interest because it allows comparisons across different morphologies and can be remotely sensed. Our results offer promise for real-time landscape-level monitoring of DIM and its global impacts in the near term.

## Introduction

Drought-induced forest mortality (DIM) is expected to increase across many regions with climate change (Dai 2013; Trenberth *et al.* 2014; Greenwood *et al.* 2017). Increases in DIM due to increased warming and hotter drought spells are expected to severely impact carbon cycles, species distributions, the economy, and global climate (Allen, Breshears & McDowell 2015). To anticipate DIM, indicators must allow real time monitoring of DIM risk. At the landscape level, such indicators must accurately predict DIM risk independently of physiological and morphological variation across and within species causing plants to differ in performance under drought (Cregg 1994). Much of the emphasis on plant drought responses and vulnerability to drought mortality has been at the species level (Bartlett, Scoffoni & Sack 2012; Choat *et al.* 2012). In contrast, variation within species, particularly species with wide distribution ranges and substantial morphological and physiological variation, is often overlooked despite the influence of such variation on drought performance (Cregg 1994; Tognetti, Michelozzi & Giovannelli 1997; Sergent, Bréda, Sanchez, Bastein & Rozenberg 2014; Garcia-Forner, Sala, Biel, Savé & Martínez-Vilalta 2016; Umaña & Swenson 2019). Hence, accurate, real time monitoring of DIM risk at large scales requires indicators of DIM risk that are robust and consistent regardless of variation in plant morphology and physiology. Additionally, for proactive management purposes, DIM indicators are most useful when, in addition to capturing the degree of DIM risk at a given level of drought, they can also distinguish the threshold at which a healthy population (no risk) becomes at risk of DIM (incipient mortality threshold: Martinez-Vilalta, Anderegg, Sapes & Sala 2019; Sapes *et al.* 2019). Thus, the first step towards accurate, real-time, and large-scale monitoring of DIM risk is identifying indicators that show both consistent incipient mortality thresholds across populations as well as high predictive power of the actual DIM risk.

Mortality thresholds and the predictive power of DIM risk indicators may vary due to variation in morphology or physiology within and across species. For instance, wood density is positively related to resistance to cavitation (Hacke & Sperry 2001; Delzon, Douthe, Sala & Cochard 2010) and denser wood has been related to lower mortality rates (Greenwood *et al.* 2017; Liang, Ye, Liu & Brodribb 2021). However, denser wood has also been shown to lead to higher mortality under drought (Hoffmann, Marchin, Abit & Lau 2011) highlighting that compensating factors such as deep roots (Padilla & Pugnaire 2016) or other strategies to avoid low water potential may offset the role of wood density. Similar patterns and offsetting mechanisms may occur within species if there is intraspecific variation in physiology and morphology. This example illustrates that robust indicators of DIM risk should be insensitive to such variation so they can be used across a broad range of plant types. Equally important is to identify indicators that can accurately assess DIM risk from any organ available. For instance, leaf-based indicators will not be useful in plants that rely on leaf shedding to survive under drought (Daubenmire 1972; Hastings, Oechel & Sionit 1989). Indicators that can be measured in any organ are advantageous because they provide greater measurement opportunities. However, to be useful for real-time, large-scale monitoring of DIM risk, organ-specific indicators must have high predictive power and be consistent among each other. Further, DIM risk indicators that integrate both morphological and physiological adjustments are of critical interest because they are likely to show low variability in mortality thresholds. Hence, assessing the relative consistency and predictive power of candidate predictors of DIM risk across populations and organs is important to identify useful large-scale indicators of DIM risk and the first step prior to assessments across species.

Given that plants combine multiple strategies to maintain water balance (Mencuccini, Minunno, Salmon, Martínez-Vilalta & Hölttä 2015), indicators of DIM that do not integrate their combined effects may not show consistent incipient mortality thresholds and predictive power among populations with varying strategies. Research on the mechanisms of DIM has identified failure of water transport -measured as percent loss of conductivity (PLC)- as a common process leading to DIM (Anderegg, Berry & Field 2012; Choat *et al.* 2012; Barigah *et al.* 2013; Hammond *et al.* 2019). Non-structural carbohydrate (NSC) availability has also been shown to be important (O’Brien, Leuzinger, Philipson, Tay & Hector 2014; Sapes *et al.* 2019), although the role is more variable and still uncertain. While both water transport and NSC are needed to maintain water balance and avoid DIM, other critical factors such as capacitance (Mcculloh, Johnson, Meinzer & Woodruff 2014) and turgor (Guadagno *et al.* 2017, but see resurrection plants) are not integrated in PLC or NSC measurements alone. Martínez-Vilalta *et al.* (2019) reasoned that water status variables would be useful indicators of DIM risk because they directly reflect the plant water balance and integrate the diverse processes and strategies -both morphological (Daubenmire 1972; Maherali & Pockman 2004; Matías, González-Díaz & Jump 2014) and physiological (Subbarao, Nam, Chauhan & Johansen 2000; Meinzer *et al.* 2016)- that plants use to avoid lethal dehydration. From all the water status variables, Martínez-Vilalta *et al.* (2019) highlighted water potential and water content (including relative water content -RWC) as potential candidates for assessment of DIM risk. However, they noted that large scale predictions based on water potential would be complicated by the enormous variability of water potential across co-occurring species (Bartlett *et al.* 2012; Choat *et al.* 2012), and suggested that RWC would be less variable and likely more integrative. Consistent with this, Sapes *et al.* 2019 showed that, in potted conditions, plant volumetric water content integrates the effects of PLC and NSC depletion - critical physiological processes leading to dehydration and DIM- and predicts DIM risk with high accuracy.

Indicators of plant water content are likely to display consistent incipient mortality thresholds and these thresholds are likely detectable via remote sensing. To remain alive and functional, cells must avoid permanent turgor loss which leads to cellular damage and plant death (Guadagno *et al.* 2017). Because cell turgor requires maintaining cell water volume above certain thresholds, plants are likely to show increasing mortality risk once RWC reaches the turgor loss point. RWC is of special interest because it reflects the amount of water present in a plant, organ, or tissue relative to the maximum it can hold (Barrs & Weatherley 1962; see below). Capturing the true maximum water content of plants can be sometimes as methodologically challenging as finding the true maximum hydraulic conductivity, the true minimum water potential in the field, or the true NSC concentrations of a species (Sanders & Arndt 2012; Quentin *et al.* 2015; Fontes & Cavender-Bares 2019). However, when maximum water content is correctly captured, RWC thresholds will likely be more consistent among individuals or species because RWC accounts for potential differences in maximum water content due to varying anatomy. Additionally, RWC reflects the integrated effects of the physiological and morphological adjustments that plants undergo to maintain water balance and prevent turgor loss, and it is likely to be more consistent among organs, populations, and -potentially- species than other less integrative indicators. Accordingly, Bartlett *et al.* 2012 showed that turgor loss occurs at similar RWC (but different water potential) across species from different biomes, which inherently vary in morphology and physiology. The common RWC values at turgor loss across species could indicate a common degree of maximum dehydration that most woody plants can withstand prior to risking death. Lastly -and unlike PLC and NSC-, a clear advantage of plant water content is that it can be estimated from organs to ecosystems via remote sensing (Ullah, Skidmore, Naeem & Schlerf 2012; Wang & Li 2012; Mirzaie *et al.* 2014; Rao, Anderegg, Sala, Martínez-Vilalta & Konings 2019; Marusig *et al.* 2020). Several remote sensing studies have observed declines in canopy water content followed by increased drought mortality (Saatchi *et al.* 2013; Asner *et al.* 2015). The potential for cross-species consistency and the fact that water content can be remotely sensed, make RWC a great candidate for large-scale assessments of DIM risk.

We performed a greenhouse drought experiment based on the point of no return (i.e., no recovery after re-watering) with one-year-old ponderosa pine (*Pinus ponderosa* Douglas ex C. Lawson) seedlings to assess the extent to which morphological and physiological variation translate into differences in incipient population-level mortality thresholds and predictive power of four potential indicators of DIM risk: PLC, NSC, water potential, and RWC. To assess the consistency and incipiency of the mortality threshold, we developed a method based on the third derivative of the mortality curve. We used a common garden approach with two genetically differentiated seedling provenances (North Plateau and Northern Rocky Mountain) (Potter, Hipkins, Mahalovich & Means 2013) known to differ in responses to drought (Cregg 1994). We focused on variability among populations and organs. Specifically, we withheld water and asked 1) do populations differ in mortality rates over time?, 2) if so, what physiological and morphological differences contribute to differences in mortality rates?, and 3) do water status variables such as water potential and RWC show high predictive power and consistent incipient mortality thresholds across populations and organs?

## Materials and Methods

### Study Design

We performed a greenhouse drought experiment at the University of Montana greenhouse facilities with one-year old ponderosa pine (*Pinus ponderosa* Douglas ex. C. Lawson) seedlings from two genetically differentiated populations known as the North Plateau race (NP) (42.6 N 122.8 W) and the Northern Rocky Mountain race (RM) (45.9 N 104.5 W) (Potter *et al.* 2013). Ponderosa pine is one of the most widely distributed species in North America and has been extensively used as a representative gymnosperm in ecological, physiological, and forestry studies. We chose one-year old seedlings because of their biological relevance for regeneration of lower tree-line forests which is constrained by dry conditions (Simeone *et al.* 2019). On May 25^th^ 2016 we planted 250 individuals from seed provided by the USDA Forest Service in 7.62 cm diameter x 43 cm tall pots using a homogeneous soil mixture consisting of 3:1:1 sand, peat moss, and top soil. Seeds started to germinate by June 2^nd^ and seedlings were grown at soil field capacity until they were big enough to be measured (ca. 6 cm height and 2.5 cm basal diameter), which corresponded to February 24^th^ 2017. Soil field capacity corresponded to soil volumetric water content values (*VWC_s_*) of ca. 20%. We monitored changes in *VWC_s_* using Meter 5TE sensors placed 10 cm above the bottom of the pots in five representative seedlings of each population. Sensors were inserted through a hole previously drilled in the side of the pots to minimize disturbance of soil structure and root system damage, which started to reach the bottom of the pot by the end of the experiment.

From February 24^th^ 2017 to May 11^th^, seedlings underwent three mild drought pre- conditioning cycles to simulate early summer conditions. During the first two cycles, we dried pots down to 50% of their field capacity (*VWC_s_* = 10%) after which we watered again to field capacity. On the last cycle, pots were dried down to 25% of their field capacity (*VWC_s_* = 5%), which corresponds to a soil water potential of −0.7 MPa based on an empirical soil characteristic curve (see below), and then watered again to field capacity. This drought preconditioning provided a more realistic response to experimental severe drought since plants were able to acclimate to increasing drought as it tends to occur in natural conditions. After the drought-preconditioning, water was withheld (final drought) in all seedlings except a control group which was kept at field capacity. Based on a preliminary drought experiment to assess symptoms of mortality (see symptoms below) as a function of soil drought and to optimize sampling times and sample size, we started measurements 29 days after the start of the drought treatment.

### Sampling procedure

We assessed the degree of drought (i.e., soil water potential), seedling physiology, and mortality risk on six weekly samplings starting on day 29 of the drought treatment. At each sampling, we measured midday VWC_s_ in five randomly chosen seedlings from each population and we used VWC_s_ to estimate soil water potential based on soil water-retention curves specific for our soil type as in Sapes *et al.* (2019). VWC_s_ sensors were installed 24h prior to measurements to reach equilibrium with soil conditions. We used the same seedlings where VWC_s_ was measured to assess mid-day leaf and stem water potential. Leaf water potential was measured in a single needle bundle using a pressure chamber (PMS Instrument Company, Corvallis, OR) following methods in (Kaufmann 1968). Stem water potentials were estimated by equilibrating the water potential of a needle bundle with the stem following methods from Begg & Turner (1970) and measuring the equilibrated bundle with the pressure chamber. We also took midday measurements of stomatal conductance and dark respiration rates per unit leaf area in each seedling using a Licor 6400 XT equipped with a 6400 RED LED chamber. CO_2_, flow, temperature, and relative humidity were set constant at 400 μmol s^−1^, 100 mol s^−1^, 25 °C, and 50%, respectively. Light conditions were set at 1,000 μmol quanta m^−2^ s^−1^ for stomatal conductance measurements and the light was turned off to measure dark respiration. We scaled up stomatal conductance to the canopy level (i.e., canopy conductance) by multiplying it by canopy leaf area (see methods S1). After this, seedlings were immediately harvested, morphology was measured (see methods S1), and seedlings were transferred to zip-lock bags with a moist paper towel in a cooler to prevent water loss (Garcia-Forner *et al.* 2016). Seedlings were then transported to the laboratory within two hours for hydraulic and water content measurements (see below). Because we could not assess mortality risk in plants that were harvested, we randomly chose a second independent subset of seedlings at each sampling event to assess mortality risk at any given point during the drought (see below). While this method is sensitive to individual variation within populations (Martinez-Vilalta *et al.* 2019), it allows us to estimate probability of mortality curves based on physiological measurements that are destructive and prevent individual-level assessments of mortality risk.

### Mortality assessment

We estimated the probability of mortality at the population level over time as in Sapes *et al.* (2019). In our study, population-level mortality is defined as the proportion of individuals from each population sampled at a given time that end up dying. At each sampling event, five groups of six seedlings (total of 30) were randomly chosen, re-watered to field capacity, and kept well-watered until September 22^nd^ to assess mortality. This method ensures accurate classification of both live and dead plants at every sampling event. Then, we classified seedlings as dead, only if their canopy and phloem were completely brown and dry after a month of re-watering and no subsequent buds appeared (Cregg 1994). We calculated mortality as the proportion of dead seedlings (out of 30) at a given sampling time. Note that in our design, physiological measurements during drought were done in individual plants, while mortality measurements were conducted at the population level. Thus, one value of probability of mortality is always associated to five individual values that reflect the variation in physiological status across the population.

### Relative Water Content

Upon arrival to the laboratory, we separated roots, stems, and needles of each seedling to measure their relative water content (RWC) based on fresh, saturated, and dry weights as: ((Fresh weight-Dry weight)/Saturated weight-Dry weight)*100 (Barrs & Weatherley 1962). Saturated weight was obtained rehydrating organs by full submersion in water for 5 hours in the dark at low temperatures (3 °C). Low temperatures prevent oversaturation artifacts that arise due to artificially low osmotic potential resulting from catabolic conversion of starch into sugars (Boyer, James, Munns, Condon & Passioura 2008). In the case of stems, we rehydrated them after hydraulic conductivity measurements (see below). We calculated whole plant RWC weighed by organ biomass fraction (proportion of each organ dry mass fraction multiplied by their respective RWC). Population-level pressure-volume curves were also built using midday leaf water potentials and the corresponding leaf RWC of each individual as in Tyree *et al.* (2002) and leaf RWC at turgor loss and water potential at turgor loss were extracted for each population.

### Stem and Root Hydraulics

#### Native hydraulic conductivity

We measured stem hydraulic conductivity and root hydraulic conductance using the gravimetric method (Sperry, Donnelly & Tyree 1988) immediately after organ fresh weight measurements. We used the same apparatus and methods described in Sapes *et al.* (2019) and a micro-flow sensor (Sensirion SLI-0430, Sensirion, Inc., Staefa ZH, Switzerland). Stem segments previously used for RWC measurements were immersed in deionized water for 20 minutes to relax xylem tensions that could artificially alter conductivity values (Trifilo, Barbera, Raimondo, Nardini & Gullo 2014). After relaxation, stems were relocated to the hydraulic apparatus and each end was recut twice at a distance of 1 mm from the tips each time (total of 2 mm per side) to remove any potential emboli resulting from previous cuts, transport, and relocation (Torres-Ruiz *et al.* 2015). Stems were then connected to the hydraulic apparatus while under water, with their terminal ends facing downstream flow. The stems were then raised out of the water and the connections were checked to ensure that there were no leaks. A solution of water with 10 mM KCl degassed at 3 kPa for at least 8 hours was used for all hydraulic measurements (Espino & Schenk 2011).

First, initial background flow was measured to account for the flow existing under no pressure, which can vary depending on the degree of dryness of the measured tissue (Hacke *et al.* 2000; Torres-Ruiz, Sperry & Fernández 2012; Blackman *et al.* 2016). Second, a small pressure gradient of 5-8 kPa was applied to run water through the stem and pressurized flow was measured. Lastly, final background flow was measured, initial and final background flows were averaged, and flow was calculated as the difference between pressurized flow and average background flow. All flows were measured upon reaching stability according to three user-adjustable criteria based on absolute changes in flow, slope of change, and standard deviation of the flow across a period of 70 seconds that are automatically calculated in the code published in Sapes *et al.* (2019). Native specific hydraulic conductivity (K) was estimated in stems as the flow divided by the pressure gradient used and standardized by xylem area and length. After conductivity measurements, stems were rehydrated to obtain saturated weights for RWC measurements. In root systems, flow was measured as above and whole root native hydraulic conductance (k) was estimated as the flow divided by the pressure gradient used and standardized by xylem area at the root collar.

#### Percent loss hydraulic conductivity

Maximum stem hydraulic conductivity (Kmax) and root hydraulic conductance (kmax) were estimated at the population level as the average stem K and root k of the pre-conditioned control measured at day 0 since the onset of the drought. Such a population approach was necessary because 1) destructive measurements in these small seedlings prevented successive measurements of K and water potential on the same individuals, and 2) flushing and vacuum infiltration techniques to obtain Kmax from embolized tissues can generate artifacts and overestimate Kmax (Cochard *et al.* 2013). Percent loss of stem conductivity and root conductance were estimated for each measured seedling as 100*(Kmax-K)/Kmax and 100*(kmax-k)/kmax, respectively. Note that negative PLC values may occur when K or k in a given sample is larger than Kmax estimated as the average K or k of controls. We calculated wood PLC weighted by organ fraction. Stem and root PLC were multiplied by their respective biomass fractions such that the whole-plant estimated value was weighted by the biomass contribution of each type of organ. Root and stem PLC can be averaged together because they are unit-less indexes that represent the relative loss of water transport capacity of their respective organs. Because we did not measure PLC in needles, wood PLC represents the overall hydraulic integrity of the stem and root systems. We excluded negative PLC values resulting from uncertainty around population level estimates of Kmax (Sapes *et al.* 2019). We ensured that exclusion of data was homogeneous across populations and sampling times (Table S1) to avoid influencing comparisons between populations.

### Non-structural Carbohydrates

After hydraulic measurements, non-structural carbohydrates were analyzed in all tissue types (needles, stems, and roots) collected at harvest. A sample of each tissue was microwaved for 180 seconds at 900 Watts in three cycles of 60 seconds to stop metabolic consumption of NSC pools, and oven-dried at 70 °C. Samples were dried to a constant mass and finely ground into a homogenous powder. Approximately 11 mg of needle tissue and 13 mg of stem or root tissue were used to analyze NSC dry mass content following the procedures and enzymatic digestion method from Hoch, Popp & Körner (2002) and Galiano, Martínez-Vilalta, Sabaté & Lloret (2012). NSC, starch, sucrose, and glucose + fructose tissue concentrations were then multiplied by their respective tissue fraction (the dry mass of a tissue relative to whole-plant dry mass) to obtain whole-plant concentrations. Whole-plant dry mass for each seedling was calculated by combining the dry mass of all samples and the remaining biomass. Finally, we calculated percent NSC relative to controls to standardize by existing differences in total NSC pools between populations at the beginning of the drought. For the sake of space, these methods are described in detail in Methods S1.

### Statistical analyses

We assessed differences in morphology between populations across the full set of measurements using two-tailed Student’s t-test for independent samples. Canopy leaf area, root to shoot ratios, organ biomass, whole-plant biomass, and plant length were used as dependent variables and population was the categorical variable in all tests. Differences in maximum stem hydraulic conductivity between populations (based on K of control plants) were also tested using the same approach. Variables were transformed to achieve normality and homogeneity of variances when needed. We also assessed potential differences in hydraulic conductivity due to plant length given that trees are known to increase hydraulic conductivity at the base of the stem as they grow tall to minimize the resistance of the hydraulic pathway (Olson *et al.* 2018). Differences in hydraulic conductivity as a function of plant length were assessed in two linear models with stem hydraulic conductivity and root hydraulic conductance as response variables, respectively, and plant length, population, and their interaction as predictors. Response variables were log-transformed to meet model assumptions.

We assessed differences in response to drought over time between populations by splitting the drought into early drought (days 0 and 29) and late drought (days 29 to 72). This was necessary because physiological measurements were not taken between days 0 and 29 to maximize sampling after the onset of mortality. Thus, analyses from day 29 to 72 reflect responses close to and after the onset of mortality in the population (when a few individuals start to die) and characterize the processes that either prevent or ultimately lead to death. Early differences in response to drought between populations were tested using two-tailed Student’s t-test for independent samples at day 0 and 29. T-tests were used instead of regressions because data included only two days. Soil and plant water potentials, canopy conductance, PLC, NSC relative to controls, and RWC across organs at day 0 and 29 were used as dependent variables while population was the categorical variable in all tests. Contrasts from day 0 to 29 provide information of whether observed differences after the onset of mortality originated during early stages prior to mortality.

Differences in response to drought between populations after day 29 were assessed using three sets of regression models with data until day 72. Response variables were i) soil, leaf, or stem water potential to represent the degree of drought intensity; ii) wood PLC, whole-plant NSC, or whole-plant RWC to represent loss of hydraulic function, NSC depletion, and degree of dehydration; and iii) population-level mortality to represent the probability of DIM. All models had days since the 29^th^ day of drought, population, and the interaction of both factors as the predictor variables. Generalized linear models (Mardia, Kent & Bibby 1979) with binomial distribution and logit link were used for models including probability of mortality. Linear models were used for all other cases as the response variables showed a linear response with time or could be transformed to meet assumptions of linearity.

We took a residuals approach to explore whether differences in dehydration rates leading to death between populations were driven by morphology, physiology, or both. First, we extracted the residuals from a model with RWC as the response variable and days since the onset of drought as the predictor. In this initial model, the residuals reflect variation in dehydration rates across populations not explained by time under drought due to individual variation in morphological and physiological characteristics. We chose RWC as our response variable because dehydration is the main process leading to DIM (Tyree *et al.* 2003; Martinez-Vilalta *et al.* 2019; Sapes *et al.* 2019) and integrates the physiological (e.g. low stomatal conductance, maintenance of hydraulic function, NSC depletion) and morphological (e.g. low canopy area, high root to shoot ratios, reduced growth) responses to prevent dehydration leading to DIM risk. Then, we built a second model for morphology with the residual variation of the first model as the response variable and population, root to shoot ratios, whole-plant biomass and all the interactions as predictive variables. In this model, significant effects of morphological variables indicate that, regardless of population of origin, morphology explains variation in dehydration rates that is not explained by time under drought. Significant interactions between a given morphological variable and population indicate that the effects of morphology are population-specific. Thus, a significant effect of population alone or as part of an interaction indicates that morphological differences alone are not able to explain all the variation in dehydration rates between populations. We followed the same approach to test the influence of physiology in a third model where we used physiological variables (stomatal conductance, dark respiration, wood PLC, and whole-plant NSC) instead of morphological variables as predictive variables. Morphological variables were log-transformed to meet model assumptions. Other physiological (e.g., soil and plant water potential) or morphological (e.g., plant length and canopy area) variables were excluded from these models because they were highly correlated with the selected predictor variables. Using whole-plant variables also allowed us to account for potential organ-specific effects without increasing the number of predictors or violating assumptions of co-linearity.

We used logistic regressions (Walker & Duncan 1967) to assess the predictive power and consistency of each mortality predictor and search for potential thresholds indicative of incipient mortality risk. For each organ within a population, logistic models included probability of mortality as the response variable and RWC or water potential as predictors. We did not include PLC and NSC relative to controls in the predictor comparison because the use of population-level reference parameters (Kmax and Control NSC) adds uncertainty that can influence predictive power, thus leading to biased comparisons. Probability of mortality was expressed as a decimal fraction following requirements of models with binomial distributions. Predictive power was estimated as the proportion of variance explained (VE) by each model (i.e., [1- residual variance/null variance] x 100) (Guisan & Zimmermann 2000). In logistic models, VE is often discouraged because it tends to underestimate predictive power due to the lack of values between 0 and 1 in the response variable. However, population-level mortality does contain intermediate values between 0 and 1 thus overcoming this issue. We assessed the consistency of mortality curves within each predictor by testing differences in slopes and intercepts of mortality relationships across populations and organs using another set of logistic models containing the interaction between a given predictor, population, and organ. In these models, non-significant interactions were not removed because our hypothesis explicitly tested the interaction between population, organ, and each predictor.

We identify incipient mortality thresholds in logistic regression models as the value of the physiological predictor beyond which mortality risk suddenly increases. To identify incipient threshold values in each mortality curve, we used a mathematical approach similar to Gregorczyk (1998) and Mischan, Pinho & Carvalho (2011) based on the third derivative of the logistic regression function (Fig. 1, see Methods S2). The third derivative describes changes in the acceleration of mortality rates as physiological status (the predictor variable) declines. It detects incipient mortality thresholds as a peak - maximum or minimum depending on the nature of the physiological predictor - near the wet end of the range of physiological values (Fig. 1), indicating a sudden change of acceleration of mortality rates (i.e., a threshold). We extracted threshold values for all mortality curves (Methods S3) and compared their degree of incipiency (probability of mortality at the threshold) and their consistency among physiological predictors. To assess consistency among organs and populations within a given predictor, we constructed a unitless index that considers the average value of the physiological predictor at the mortality threshold across all mortality curves and describes how the threshold of each curve deviates from the average value (Methods S2). We compared threshold incipiency, threshold consistency, and variance explained among predictors using ANOVA and Tukey Honest Significant Difference tests. All analyses were performed using R version 3.6.3 (R Core Team 2020).

**Figure 1:**
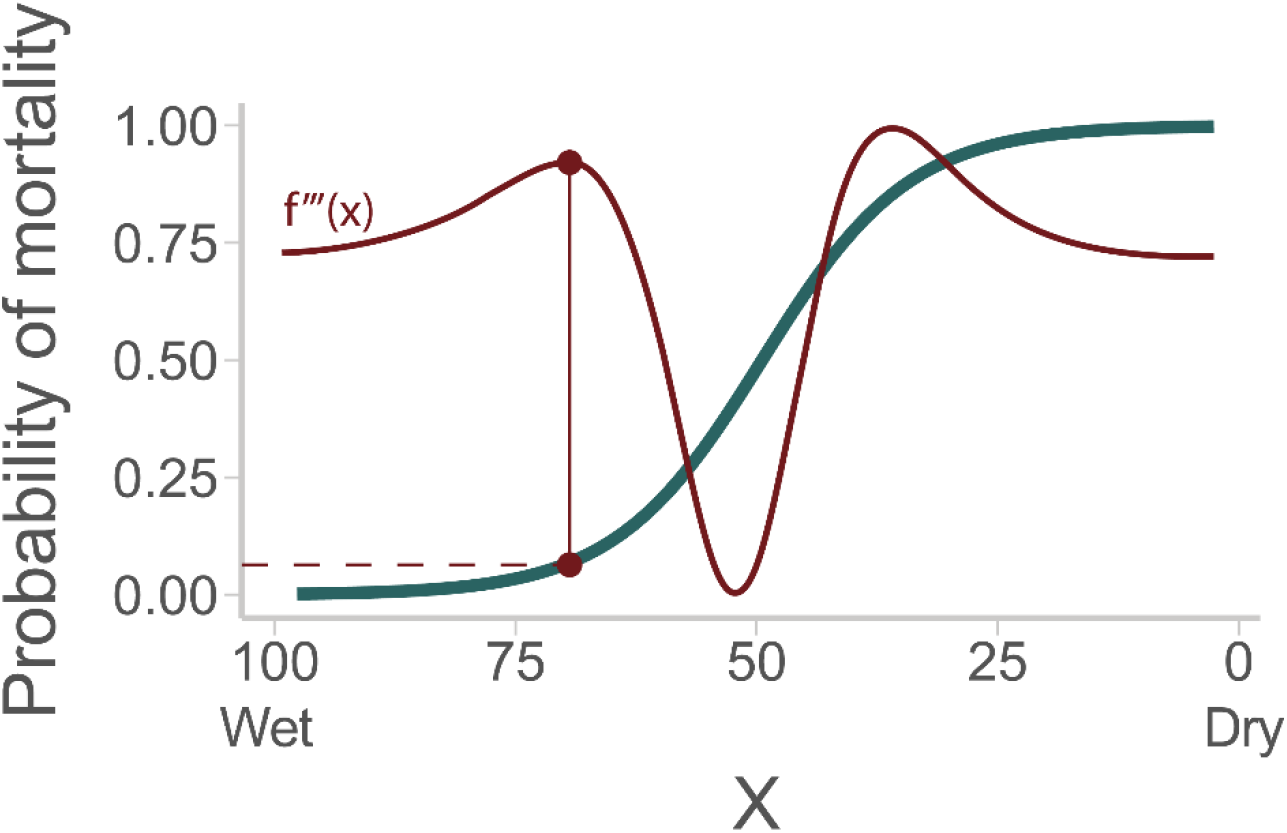
The third derivative (red curve) identifies the point of the mortality curve (teal curve) at which mortality risk starts to suddenly increase (incipient mortality threshold) in response to physiological decline. The incipient mortality threshold (red dot) appears in the third derivative as a peak -maximum or minimum depending on the nature of the physiological predictor- near the wet end of the range of physiological values. Once the incipient mortality threshold is identified, we can evaluate the probability of mortality at which such threshold occurs (red dashed line) and determine the degree of incipiency. The physiological value (X) of the incipient threshold can vary among mortality curves of different organs and populations. Little variation among curves leads to more consistent thresholds.

## Results

North Plateau (NP) seedlings were longer (from root tip to stem tip) (t = 2.51, p = 0.014; Fig. 2a) and had greater whole-plant biomass (t = 3.58, p < 0.001; Fig. 2b) than Northern Rocky Mountain (RM) seedlings (all dates combined). These differences were due to greater stem (t = 5.04, p < 0.001; Fig. S1a) and root biomass (t = 5.43, p < 0.001; Fig. S1b) in NP seedlings. Differences in biomass allocation translated to greater root to shoot ratios in NP seedlings (t = 4.83, p < 0.001; Fig. 2c). Differences in plant length between populations were associated with greater hydraulic conductivity in NP seedlings (stem: R^2^_adj_ = 0.18, plant length: p < 0.001, population: p = 0.077; roots: R^2^_adj_ = 0.08, plant length: p = 0.003, population: p = 0.047; Table S2). Accordingly, NP seedlings also had greater stem maximum hydraulic conductivity (t = 1.94, p = 0.059; Fig. S1c).

**Figure 2.**
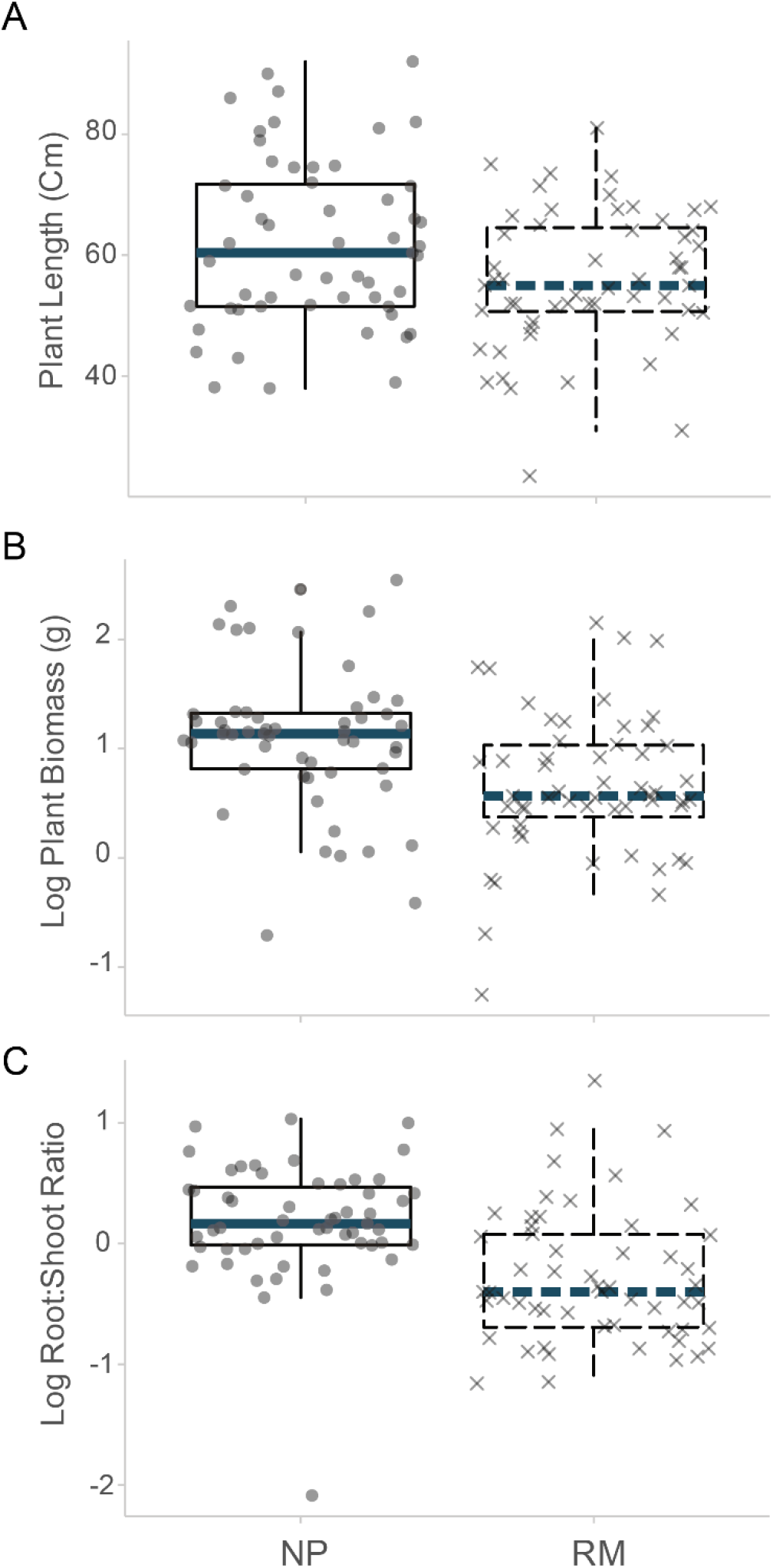
Morphological differences between North Plateau (NP) and Rocky Mountain (RM) seedlings. NP seedlings were larger in size and biomass and allocated greater biomass to below ground organs. Differences among populations are significant across all panels.

Up to day 29, the probability of mortality was zero in both populations (Fig. 3e). Mortality risk started to increase above zero by days 29 and 42 in NP and RM populations, respectively (Fig. 3d). After the onset of mortality in each population, mortality probabilities increased at the same rate in both populations.

**Figure 3.**
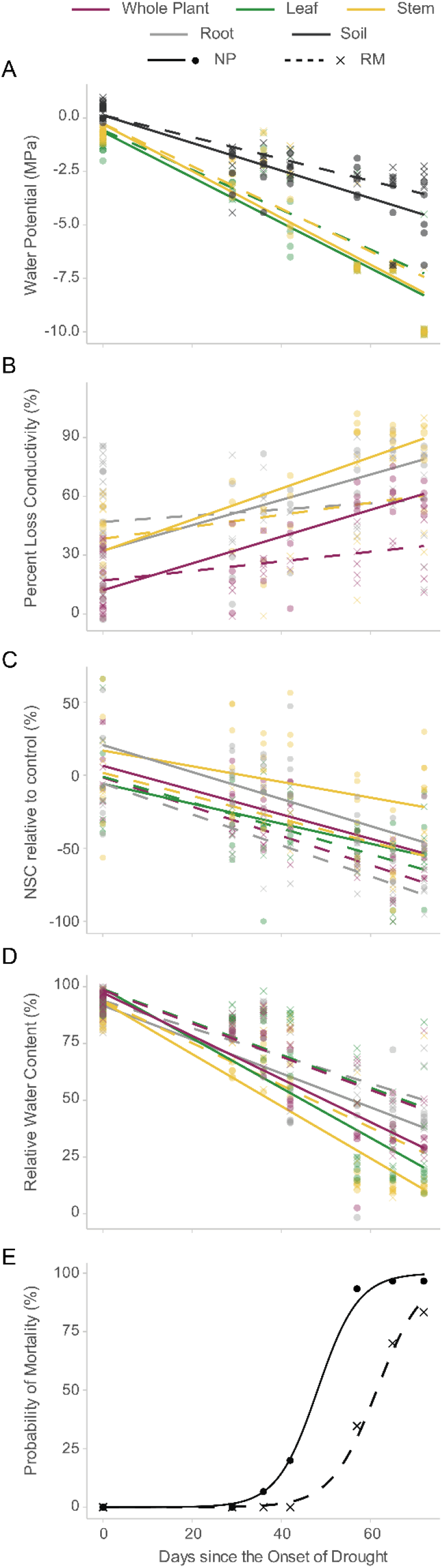
Changes in population-level mortality, drought intensity, and whole-plant physiological status over time. Panel A: Water potentials decreased over time in soil (black), stem (yellow) and leaves (green) but North Plateau (NP, circles and corresponding solid line) seedlings experienced greater decline rates than Rocky Mountain (RM, crosses and corresponding dashed line) seedlings. Panel B: Both populations experienced loss of conductivity over time in both stems and roots (gray) and at the whole-plant level (purple), but NP seedlings lost hydraulic conductivity at faster rates in all organs. Panel C: Non-structural carbohydrates declined over time in both populations, but NP seedlings experienced greater root NSC depletion rates from day 29 onward. Panel D: Relative water content declined over time at similar rates across all organs, but NP seedlings dehydrated faster than RM seedlings. Panel E: Probability of mortality increased after day 29 and 42 of drought in NP and RM seedlings, respectively.

At the beginning of the experiment (day 0), populations only showed physiological differences in dark respiration, with RM seedlings respiring more than NP seedlings (t = 2.25, p = 0.04; data not shown). These differences disappeared by day 29 of drought. In contrast, at day 29, NP seedlings had lower whole-plant RWC (t = −4.57, p = 0.002; compare solid and dashed purple lines), which was driven by lower stem (yellow) and root (gray) RWC (stem: t = −4.70, p = 0.003; roots: t = −2.25 p = 0.056; Fig. 3d). These results indicate that NP seedlings experienced greater dehydration rates during the early stages of the drought.

At late stages of drought (day 29 to 72), the soil water potentials of the two populations clearly diverged (compare solid and dashed black lines in Fig. 3a) and continued to decrease at similar rates (R^2^_adj_ = 0.30, days: p < 0.001; population: p = 0.029; Table S3). The same pattern was observed for leaf (green) and stem (yellow) water potentials (needles: R^2^_adj_ = 0.77, days: p < 0.001, population: p = 0.031; stems: R^2^_adj_ = 0.83, days: p < 0.001, population: p = 0.065; Fig. 3a, Table S3). Declines in water potential were accompanied by increases in PLC in both populations, but PLC in NP seedlings increased faster than in RM seedlings (R^2^_adj_ = 0.57, days: p < 0.001, days x population: p < 0.022; Fig. 3b, Table S3). This pattern was driven by higher PLC rates in roots (R^2^_adj_ = 0.27, days: p < 0.001, population: p = 0.044, days x population: p < 0.011; gray solid vs. dashed lines in Fig. 3b, Table S3), which were explained by higher root to shoot ratios in NP seedlings (R^2^_adj_ = 0.60, days: p < 0.001, root shoot ratio: p = 0.035, days x population: p = 0.018; Fig. S2, Table S3). By day 29, NP seedlings also had higher stem (yellow) and root (gray) NSC (relative to controls) than RM seedlings. However, they also consumed NSC in their roots at a faster rate than RM seedlings throughout the drought (plant: R^2^_adj_ = 0.57, days: p < 0.001, population: p < 0.001; needles: R^2^_adj_ = 0.16, days: p < 0.001; stems: R^2^_adj_ = 0.38, days: p < 0.001, population: p < 0.001; roots: R^2^_adj_ = 0.16, days: p < 0.001, days x population: p = 0.014; Fig. 3c, Table S3). Finally, NP seedlings lost RWC in needles at faster rates than RM seedlings (solid vs dashed lines in Fig. 3d; R^2^_adj_ = 0.78, days: p < 0.001, days x population: p = 0.030; Table S3). In turn, differences in dehydration rates in needles led to marginally lower canopy conductance rates in NP seedlings (R^2^_adj_ = 0.03, days: p = 0.071, population: p = 0.055, days x population: p = 0.079; Table S3). While leaves of NP seedlings dehydrated faster (i.e., different slopes between populations; solid vs dashed green lines in Fig. 3d), there were no population differences in dehydration rates at the whole plant level (similar slope of solid and dashed purple lines) because needles were a very small fraction of the biomass and therefore contribute little to the whole-plant water content. However, whole-plant RWC still differed between populations (i.e., different intercepts between populations) (R^2^_adj_ = 0.71, days: p < 0.001, population: p < 0.001; Fig. 3d, Table S3) as observed during early drought stages (day 29). As a result, NP seedlings started dying (the point at which mortality risk started to increase) more than two weeks earlier but both populations showed similar rates of mortality once mortality started (days: p < 0.001; population: p = 0.024; Fig. 3e, Table S3).

Residual analyses allowed us to determine whether differences in dehydration rates between populations were explained by differences in morphology, physiology, or both. Variation in dehydration rates unrelated to time since drought was partially attributed to physiology but population was still significant (R^2^_adj_ = 0.46, population: p = 0.05, respiration: p = 0.005, population x stomatal conductance: p = 0.002, respiration x wood PLC: p = 0.001, population x stomatal conductance x wood PLC: p < 0.001). This indicates that physiological differences alone cannot fully explain the differences in dehydration rates observed between populations. On the other hand, population was not significant when the remaining variation was attributed to morphology (R^2^_adj_ = 0.32, log(root to shoot ratio): p = 0.002, log(plant biomass): p = 0.013). This indicates that morphological differences in root to shoot ratios and plant size alone can explain the differences in dehydration rates observed between populations. That is, morphological variables absorb the variation otherwise explained by the categorical variable ‘population’ thus making it not significant.

The ability of water potential, and RWC to predict mortality was similar among populations and organs, both in terms of predictive power and degree of significance (Figs. 4c, 5; Table S5). Water status variables (RWC and water potential) had comparably high predictive power (RWC: p-value_Range_ = <0.001 - 0.002, VE_Average_ = 76.35%, VE_Range_: 54%-95%; water potential: p-value_Range_ = <0.001 - 0.01, VE_Average_ = 74.42%, VE_Range_: 46%-93%). Additionally, both variables showed similar relationships with DIM risk across organs and populations as supported by the lack of significant differences in intercepts and slopes (Fig. 5a-b, Table S6). Logistic regressions identified mortality thresholds in both variables (Fig. 5). RWC and water potential showed incipient mortality thresholds in all organs measured and in the two populations (Fig. 5a-b). RWC showed the greatest degree of incipiency in mortality thresholds (Fig. 4a). Importantly, both RWC and water potential showed mortality thresholds close to the values corresponding to leaf turgor loss (Fig. 5a-b, vertical lines). Note, however, that the high proximity of the incipient mortality threshold to the turgor loss point could be due to variability among individuals, which could shift the incipient mortality threshold in population-level mortality curves (Martinez-Vilalta *et al.* 2019). If so, the onset of mortality risk at the individual level might be slightly further away from turgor loss (Mantova, Menezes-Silva, Badel, Cochard & Torres-Ruiz 2021).

**Figure 4.**
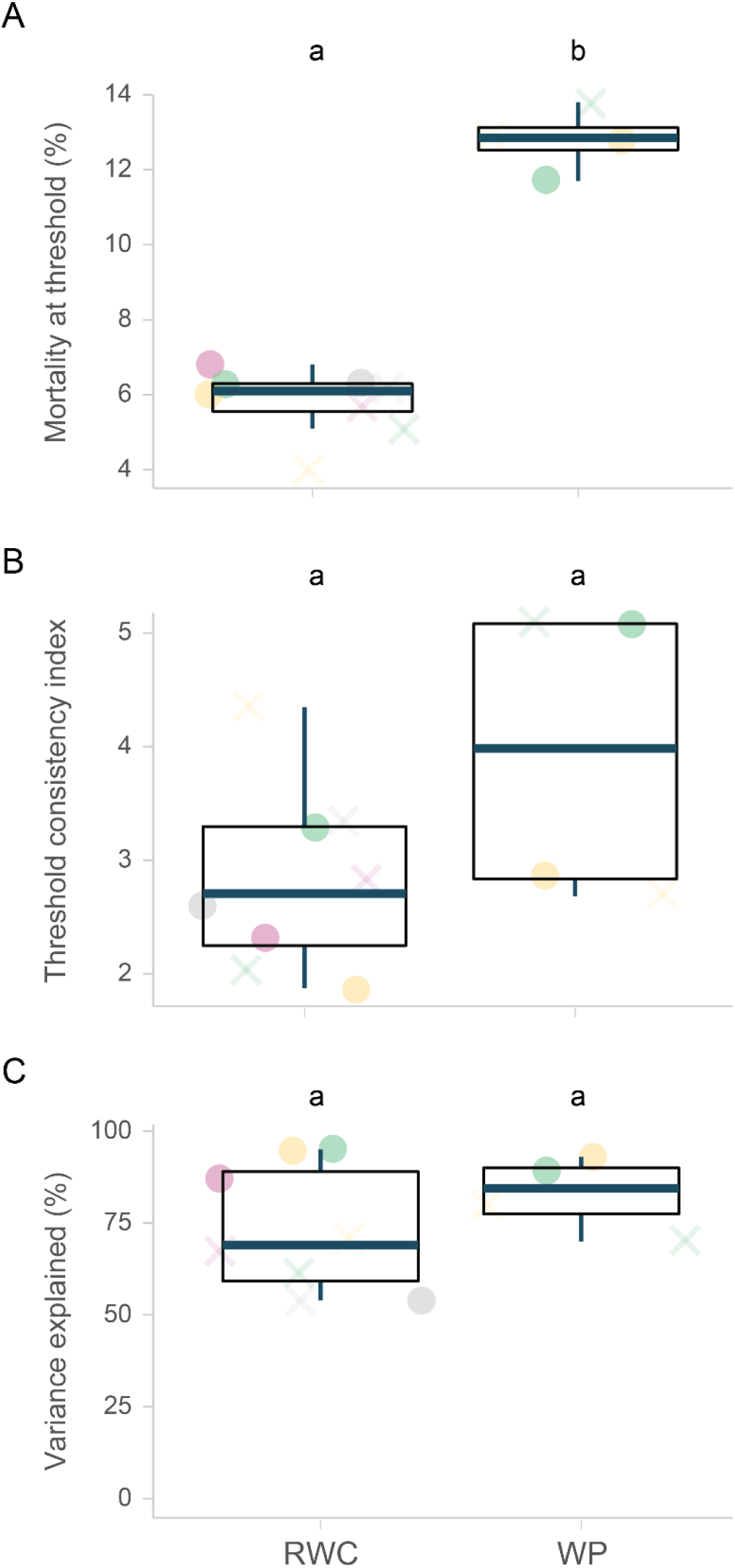
Degree of threshold incipiency, threshold consistency, and predictive power of predictors of mortality associated with water status. Relative water content (RWC) and water potential predicted DIM risk in all organs (gray: roots, yellow: stems, green: leaves, and purple: plant) and populations (circles: North Plateau (NP), crosses: Rocky Mountain (RM)) and had a similar average predictive power (i.e., variance explained), variability in predictive power, and threshold consistency. However, RWC showed the most incipient thresholds.

**Figure 5.**
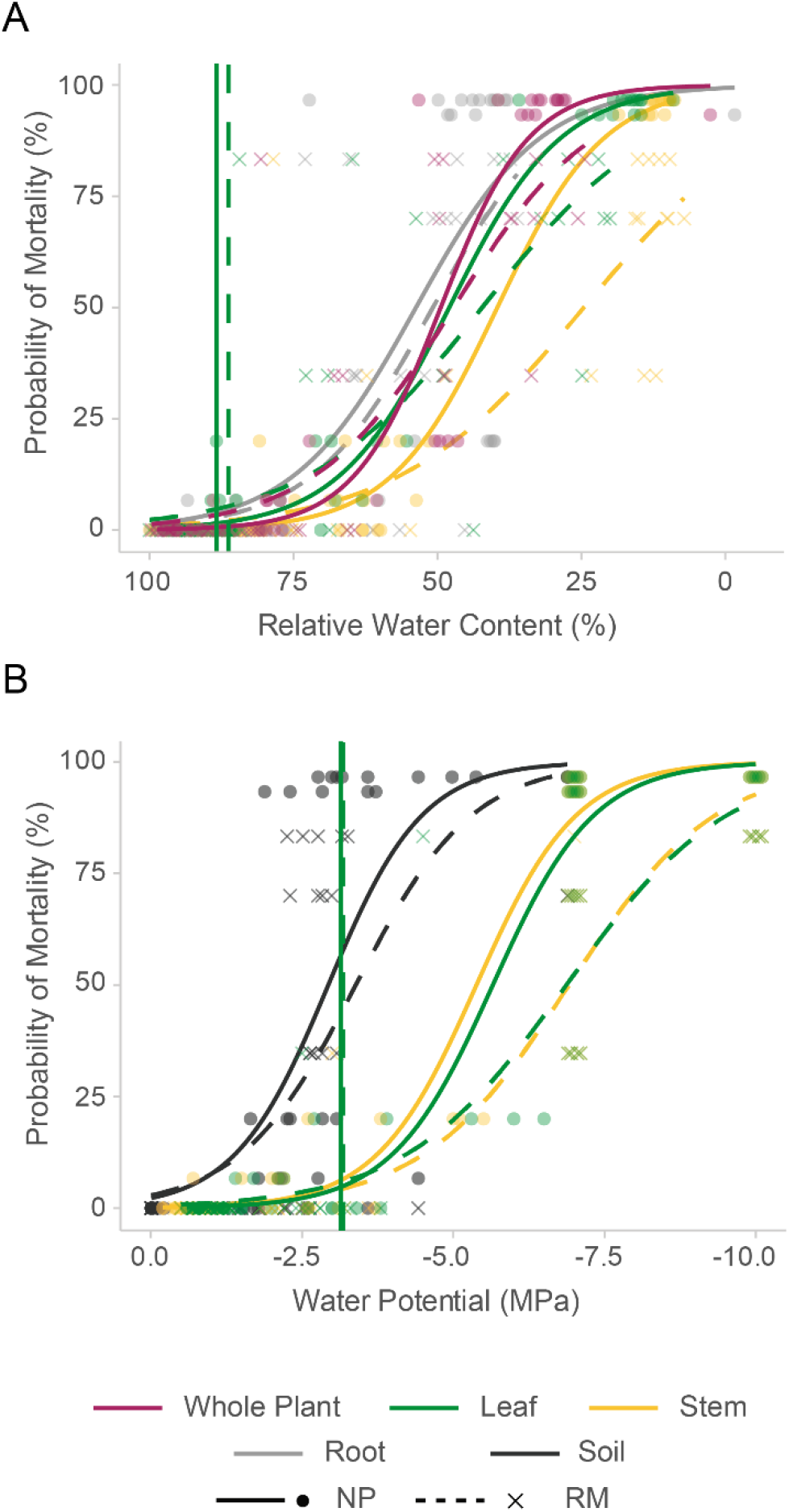
Relationships between DIM risk and DIM predictors associated with water status. Both relative water content (RWC) and water potential showed similar mortality functions with incipient mortality thresholds after turgor loss (vertical lines) across leaves (green), stems (yellow), roots (gray), and whole plant level (purple) and populations of NP (circles and corresponding solid lines) and RM seedlings (crosses and corresponding dashed lines). Note that black water potential curves correspond to soil water potential.

## Discussion

It has long been known that individuals of the same species can show morphological and physiological differences across the landscape. Our results in a ponderosa pine common garden experiment suggest that such differences can affect the predictive power and mortality thresholds of DIM risk indicators. Hence, it is critical to find which DIM indicators are the least sensitive to individual variation across the landscape. Our results show that DIM indicators related to plant water status show high and consistent predictive power across populations and organs. Among these, RWC standardizes morphological differences among individuals, likely explaining why it provided consistent incipient mortality thresholds among populations and organs. Because of its consistency and high predictive power -and because it can be estimated via remote sensing-, RWC could be a promising indicator to monitor real time DIM risk across the landscape.

Morphological and physiological differences between populations played in concert under drought stress and explained dehydration rates and time to death. All pots were similar in size and contained the same soil mixture thus holding similar amounts of water at the start of the drought. Hence, differences in dehydration rates between populations can be attributed to differences in water consumption during the drought. NP seedlings grew taller and produced more biomass than RM seedlings and had built much greater root biomass prior to the onset of drought resulting in greater root:shoot ratios (Fig. 2a-c). NP seedlings also grew longer from the tip of roots to the tip of shoots prior to the drought (Fig. 2a), which, in the absence of compensating factors, would increase the resistance to water flow through the plant due to a longer hydraulic pathway. Perhaps to compensate for this increase, NP seedlings developed greater maximum stem hydraulic conductivity, a response that was related to plant length (Fig. S1, Table S2; Olson *et al.* 2018). Such pattern is caused by downwards-widening of stem xylem conduits and is likely to extend into the roots (Carrer, Von Arx, Castagneri & Petit 2014; Olson *et al.* 2018). However, plants can also reduce resistance through other mechanisms such as more efficient -but vulnerable- pits (Delzon *et al.* 2010; Roskilly, Keeling, Hood, Giuggiola & Sala 2019). Indeed, population differences in maximum hydraulic conductivity could not be fully explained by plant length (significant population effect in root hydraulic conductance, Table S2) suggest that other interacting mechanisms are at play. Increased stem hydraulic conductivity along with greater root biomass allowed NP seedlings to transport more water (i.e., greater hydraulic efficiency) and maintain growth rates under well-watered conditions. However, greater hydraulic efficiency and plant size came at the cost of greater soil water consumption (Fig. 3a), lower soil water potential, greater cavitation rates (Fig. 3b, Fig. S2) and earlier death during drought (Fig. 3e). The smaller size and lower hydraulic efficiency of RM seedlings was an advantage under drought and led to slower dehydration and mortality rates (Fig. 3d-e). RWC captured all this morphological and physiological variation and its consequences during the dehydration process: NP seedlings dehydrated faster than RM seedlings and reached lethal thresholds earlier, thus, condensing the RWC-mortality functions of both populations into a smaller space (Fig. 5a). We note that population level estimates of mortality based on subsampling are sensitive to individual variation within each population (Martinez-Vilalta *et al.* 2019). However, individual variation was similar among the two populations thus reducing bias.

The third-derivative approach we used provides a robust and objective way to identify incipient mortality thresholds. Our goal was to find indicators of DIM risk sensitive enough to detect the limit of dehydration after which mortality begins to occur. Therefore, ideal indicators should characterize well the transition from absent or very low mortality values that remain stable as water status declines, to the point when mortality starts to increase rapidly with further dehydration. We refer to this transition as the incipient mortality threshold: the point at which mortality risk starts to rapidly increase. Ideally, to capture all mortality, this point should be as close to zero mortality risk as possible. Derivatives of the mortality function can identify thresholds because their maxima and minima describe key transition points in the mortality function. The minimum of the first derivative - the point at which the monotonic increase in mortality risk starts to slow down - coincides with 50% mortality risk. Based on our definition, a 50% mortality risk is far from being incipient mortality since half of the trees would already be doomed to die. The maximum of the second derivative -the point of maximum acceleration of the mortality rate-coincides with a mortality risk value still substantially above zero (because it describes maximum acceleration, not when the acceleration starts to rapidly increase) and thus, does not identify the physiological point that precedes imminent tree death. However, the maximum of the third derivative closest to the wet end of the physiological range describes the point at which acceleration suddenly increases, which defines the onset of rapid mortality risk increase. As such, the third derivative provides the desired mortality threshold value. Third-derivative approaches have been used in the plant growth literature to identify growth phases, optimize, and predict growth (Gregorczyk 1998; Mischan *et al.* 2011). Applying derivative approaches in the field of DIM has both conceptual and practical advantages. Conceptually, it provides an objective and biologically meaningful way to determine transition points between mortality stages in trees among predictors of DIM. Practically, once mortality stages are described for a given predictor, forest managers can identify forested areas where no risk, incipient risk, high risk, and total death is expected based on the physiological status of trees within the area. Such an approach could facilitate decisions of which -or if- interventions are worth pursuing based on the current predicted mortality stage.

Large scale monitoring of mortality risk requires variables applicable at different scales and that are reasonably consistent across plant organs, populations, and species. Earlier, we argued that RWC could be a promising candidate because it reflects the interactions between water transport and carbohydrate availability that determine plant water balance (uptake and loss) (Sapes *et al.* 2019) as well as the combined effects of morphology and physiology on dehydration time (Table S4) (Martinez-Vilalta *et al.* 2019). RWC, like water potential, can also detect the transition to permanent turgor loss (using RWC_TLP_), which has been associated to cellular and organ death under drought (Guadagno *et al.* 2017). Our results across two differentiated ponderosa pine populations partially support our prediction that RWC would be the most suitable predictor of DIM risk for large scale assessments in several ways. First, RWC accurately predicted DIM risk across populations and organs despite the differences in physiology and morphology and the differences in time to death. Second, populations and organs dehydrated at different rates (Fig. 3d) but both populations started to die at the same RWC and at similar rates with decreasing RWC, and incipient mortality thresholds of RWC were also relatively consistent across organs despite known potential undersaturation methodological challenges associated with measuring RWC after embolism and cellular damage occurs. In nature, canopy dieback prior to mortality may also reflect different dehydration rates depending on hydraulic properties and canopy position, but actual organ death could still occur at similar RWC. Finally, mortality increased as RWC declined beyond RWC_TLP_, consistent with the increase in cellular damage that follows turgor loss (Guadagno *et al.* 2017). However, in contrast to our prediction, water potential was also a particularly good predictor of DIM risk, was quite consistent across organs and populations and showed consistent mortality thresholds after turgor loss point (although slightly farther from incipient mortality relative to RWC). Therefore, in ponderosa pine, both RWC and water potential are good indicators of DIM risk.

In our experiment, plant water potential also showed high predictive power and consistent mortality thresholds associated to turgor loss across both populations and organs. Low water potentials cause embolism and stomatal closure which impair water transport and carbohydrate availability, respectively (Tyree & Sperry 1989; Sevanto, McDowell, Dickman, Pangle & Pockman 2014; Meinzer *et al.* 2016). Because both reduced water transport and carbohydrate availability are involved in drought mortality (Mitchell *et al.* 2013), the predictive power of water potential is likely high (Fig. 4b). Additionally, plant water potential also indicates wilting via the turgor loss point (⍰_TLP_), thus explaining the observed degree of incipiency in mortality thresholds close to turgor loss (Fig. 5b). Indeed, these characteristics have made water potential one of the main components of mechanistic DIM models (Sperry & Love 2015). The consistent water potential thresholds and high predictive power we found in both populations suggest that -at least in ponderosa pine- there is little variability in lethal water potentials among populations. These results are consistent with relatively high vulnerability to embolism observed in ponderosa pine (Piñol & Sala 2000; Martinez-Vilalta, Sala & Pinol 2004) and low intraspecific variability in vulnerability to embolism (Maherali & DeLucia 2000; Stout & Sala 2003). However, while a good indicator of real-time DIM risk within species, plant water potential may pose challenges in landscape-level assessments when multiple species are present. The large variation in minimum water potentials (Choat *et al.* 2012) and in ⍰_TLP_ (Bartlett *et al.* 2012) across species and biomes is likely to result in different incipient mortality thresholds across species. Thus, despite its usefulness in mechanistic approaches, large-scale monitoring of DIM based on remotely sensed water potential might only be applicable in monospecific or low diversity forests in which pixels mostly contain a single species and the species identity is known. In these instances, knowledge of the ⍰_TLP_ of the species present may still be necessary to accurately predict thresholds leading to DIM within the area of interest.

Of the two indicators of water status that consistently predicted DIM between populations, RWC has important benefits compared to water potential due to its relative nature. Relative water content conveniently standardizes differences in maximum water content among organs, individuals, and -to an extent- species, thus reducing the degree of variation in dehydration leading to turgor loss point across plants (Bartlett *et al.* 2012). While assessments of the consistency of RWC thresholds across species and under different field conditions are still lacking, RWC showed consistent mortality thresholds and relationships across organs and populations. Hence, RWC can be used to monitor DIM risk in real time from any organ available such as stems or roots, which is of critical importance given that drought-deciduous species shed leaves (Daubenmire 1972). In the case of roots, we cannot estimate root water content remotely. However, and consistent with recent studies (Nardini *et al.* 2020), the predictive power of root RWC highlights the importance of root functioning and integrity during drought. Both RWC and water potential can be remotely sensed (Cohen, Alchanatis, Meron, Saranga & Tsipris 2005; Zhang, Zhou, Gentine & Xiao 2019). However, remote sensing estimates of plant water potential (i.e., xylem tension) are more likely to reflect the diversity of morphology, physiology, and drought strategies (position within the iso-anisohydric spectrum) within a pixel than to reflect DIM risk (Martinez-Vilalta *et al.* 2019). On the other hand, RWC is directly related to plant water content and can be estimated across the landscape using hyperspectral and thermal techniques (Ceccato, Flasse, Tarantola, Jacquemoud & Grégoire 2001; Elsayed, Darwish, Elsayed & Darwish 2017; Konings *et al.* 2017). Remote sensing estimates of plant water content -once standardized by the maximum values observed within a pixel- are likely to show similar properties than empirical RWC and to be a more consistent indicator across plants. While standardizing remote sensing estimates of absolute plant water content is not a task lacking in challenges, recent work has shown that this approach holds promise (Rao *et al.* 2019; Marusig *et al.* 2020). In summary, RWC stands out as a good candidate for large-scale, real-time assessments of DIM risk.

Although patterns of drought mortality inferred from potted seedlings in the greenhouse are likely different from those in the field (particularly in mature trees), our study highlights a biologically meaningful and critical point: morphological and physiological differences among populations can significantly influence mortality rates and the predictive capacity of DIM risk indicators that are not associated with plant water pools. Hence, large-scale indicators of DIM risk should be chosen based on the consistency and incipiency of their mortality thresholds and predictive power among populations. Doing so will ensure that such indicators can predict DIM risk across the landscape regardless of the existing intraspecific variation. Based on our results, RWC may achieve this because it integrates the physiological mechanisms of mortality under drought and standardizes morphological differences among individuals and populations. Future research should assess the consistency and incipiency of mortality thresholds and the predictive power of RWC across a diverse range of species and in field conditions. It is particularly critical to compare species with varying leaf types (needle and broadleaf), drought strategies (deciduous, evergreen, and resprouting), sizes, and developmental stages. Similarly, remote sensing estimates of RWC should be further developed (but see Rao *et al.* 2019; Marusig *et al.* 2020). Despite, the work that still lays ahead, we hope that our findings help increase the accuracy of current monitoring efforts and may open a path of research towards global scale assessments of DIM risk (Martinez-Vilalta *et al.* 2019).

## Acknowledgements

The authors thank L. Fishman for providing invaluable greenhouse space and constant insights and P. Demaree for his help collecting data. This project was part of a larger collaboration with S. Dobrowski and M. Maneta, who provided valuable input. A. Woods, B. Roskilly, R. Montgomery, S. Kothari, A. Castillo Castillo, R. Partelli, and L. Schroeder provided comments in early versions of this manuscript.

This work was supported by a National Science Foundation grant to AS (BCS 1461576). GS received funding from the NSF Experimental Program to Stimulate Competitive Research (EPSCoR) Track-1 EPS-1101342 (INSTEP 3). The authors thank L. Fishman for providing invaluable greenhouse space and constant insights, P. Demaree for his help collecting data and S. Dobrowski, A. Woods, B. Roskilly, R. Montgomery, S. Kothari, A. Castillo Castillo, R. Partelli, and L. Schroeder for comments in early versions of this manuscript.

## Author Contribution

G.S. and A.S. designed the experiment. G.S. collected and analyzed the data with contributions from A.S. G.S. and A.S. wrote the manuscript.

## References

Allen C.D., Breshears D.D. & McDowell N.G. (2015) On underestimation of global vulnerability to tree mortality and forest die-off from hotter drought in the Anthropocene. Ecosphere 6, 1–55.

Anderegg W.R.L., Berry J.A. & Field C.B. (2012) Linking definitions, mechanisms, and modeling of drought-induced tree death. Trends in Plant Science 17, 693–700.

Asner G.P., Brodrick P.G., Anderson C.B., Vaughn N., Knapp D.E. & Martin R.E. (2015) Progressive forest canopy water loss during the 2012–2015 California drought. Proceedings of the National Academy of Sciences 2015, 201523397.

Barigah T.S., Charrier O., Douris M., Bonhomme M., Herbette S., Améglio T., … Cochard H. (2013) Water stress-induced xylem hydraulic failure is a causal factor of tree mortality in beech and poplar. Annals of Botany 112, 1431–1437.

Barrs H.D. & Weatherley P.E. (1962) A Re-Examination of the Relative Turgidity Technique for Estimating Water Deficits in Leaves. Australian Journal of Biological Sciences 15, 413–428.

Bartlett M.K., Scoffoni C. & Sack L. (2012) The determinants of leaf turgor loss point and prediction of drought tolerance of species and biomes: a global meta-analysis. Ecology Letters 15, 393–405.

Begg J.E. & Turner N.C. (1970) Water Potential Gradients in Field Tobacco. Plant Physiology 46, 343–346.

Blackman C.J., Pfautsch S., Choat B., Delzon S., Gleason S.M. & Duursma R.A. (2016) Toward an index of desiccation time to tree mortality under drought. Plant Cell and Environment 39, 2342–2345.

Boyer J.S., James R.A., Munns R., Condon T. & Passioura J.B. (2008) Osmotic adjustment leads to anomalously low estimates of relative water content in wheat and barley. Functional Plant Biology 35, 1172–1182.

Carrer M., Von Arx G., Castagneri D. & Petit G. (2014) Distilling allometric and environmental information from time series of conduit size: The standardization issue and its relationship to tree hydraulic architecture. Tree Physiology 35, 27–33.

Ceccato P., Flasse S., Tarantola S., Jacquemoud S. & Grégoire J.M. (2001) Detecting vegetation leaf water content using reflectance in the optical domain. Remote Sensing of Environment 77, 22–33.

Choat B., Jansen S., Brodribb T.J., Cochard H., Delzon S., Bhaskar R., … Zanne A.E. (2012) Global convergence in the vulnerability of forests to drought. Nature 491, 752–755.

Cochard H., Badel E., Herbette S., Delzon S., Choat B. & Jansen S. (2013) Methods for measuring plant vulnerability to cavitation: a critical review. Journal of Experimental Botany 64, 4779–4791.

Cohen Y., Alchanatis V., Meron M., Saranga Y. & Tsipris J. (2005) Estimation of leaf water potential by thermal imagery and spatial analysis. Journal of Experimental Botany 56, 1843–1852.

Cregg B.M. (1994) Carbon allocation, gas exchange, and needle morphology of Pinus ponderosa genotypes known to differ in growth and survival under imposed drought. Tree Physiology 14, 883–898.

Dai A. (2013) Increasing drought under global warming in observations and models. Nature Climate Change 3, 52–58.

Daubenmire R.F. (1972) Phenology and other characteristics of tropical semi-deciduous forest in north-western Costa Rica. The Journal of Ecology 60, 147–170.

Delzon S., Douthe C., Sala A. & Cochard H. (2010) Mechanism of water-stress induced cavitation in conifers: bordered pit structure and function support the hypothesis of seal capillary-seeding. Plant, Cell & Environment 33, 2101–2111.

Elsayed S., Darwish W., Elsayed S. & Darwish W. (2017) Hyperspectral remote sensing to assess water status, biomass and yield ofmaize cultivars under salinity and water stress. 76, 62–7262.

Espino S. & Schenk H.J. (2011) Mind the bubbles: Achieving stable measurements of maximum hydraulic conductivity through woody plant samples. Journal of Experimental Botany 62, 1119–1132.

Fontes C.G. & Cavender-Bares J. (2019) Toward an integrated view of the “elephant”: Unlocking the mysteries of water transport and xylem vulnerability in oaks. Tree Physiology.

Galiano L., Martínez-Vilalta J., Sabaté S. & Lloret F. (2012) Determinants of drought effects on crown condition and their relationship with depletion of carbon reserves in a Mediterranean holm oak forest. Tree Physiology 32, 478–489.

Garcia-Forner N., Sala A., Biel C., Savé R. & Martínez-Vilalta J. (2016) Individual traits as determinants of time to death under extreme drought in Pinus sylvestris L. Tree Physiology 36, 1196–1209.

Greenwood S., Ruiz-Benito P., Martínez-Vilalta J., Lloret F., Kitzberger T., Allen C.D., … Jump A.S. (2017) Tree mortality across biomes is promoted by drought intensity, lower wood density and higher specific leaf area. Ecology Letters 20, 539–553.

Gregorczyk A. (1998) Richards plant growth model. Journal of Agronomy and Crop Science 181, 243–247.

Guadagno C.R., Ewers B.E., Speckman H.N., Aston T.L., Huhn B.J., DeVore S.B., … Weinig C. (2017) Dead or alive? Using membrane failure and chlorophyll fluorescence to predict mortality from drought. Plant Physiology 175, 223–234.

Guisan A. & Zimmermann N.E. (2000) Predictive habitat distribution models in ecology. Ecological Modelling.

Hacke U.G. & Sperry J.S. (2001) Functional and ecological xylem anatomy. Perspectives in Plant Ecology, Evolution and Systematics 4, 97–115.

Hacke U.G., Sperry J.S., Ewers B.E., Ellsworth D.S., Schäfer K.V.R. & Oren R. (2000) Influence of soil porosity on water use in Pinus taeda. Oecologia 124, 495–505.

Hammond W.M., Adams H.D., Yu K., Wilson L.A., Will R.E. & Anderegg W.R.L. (2019) Dead or dying? Quantifying the point of no return from hydraulic failure in drought-induced tree mortality. New Phytologist 223, 1834–1843.

Hastings S., Oechel W. & Sionit N. (1989) Water relations and photosynthesis of chaparral resprouts and seedlings following fire and hand clearing. (ed K. Sc), Natural History Museum of Los Angeles County, Los Angeles.

Hoch G., Popp M. & Körner C. (2002) Altitudinal increase of mobile carbon pools in Pinus cembra suggests sink limitation of growth at the Swiss treeline. Oikos 98, 361–374.

Hoffmann W. a., Marchin R.M., Abit P. & Lau on L. (2011) Hydraulic failure and tree dieback are associated with high wood density in a temperate forest under extreme drought. Global Change Biology 17, 2731–2742.

Kaufmann M. (1968) Evaluation of the Pressure Chamber Technique for Estimating Plant Water Potential of Forest Tree Species. Forest Science 14, 369–374.

Konings A.G., Yu Y., Xu L., Yang Y., Schimel D.S. & Saatchi S.S. (2017) Active microwave observations of diurnal and seasonal variations of canopy water content across the humid African tropical forests. Geophysical Research Letters 44, 2290–2299.

Liang X., Ye Q., Liu H. & Brodribb T.J. (2021) Wood density predicts mortality threshold for diverse trees. New Phytologist 229, 3053–3057.

Maherali H. & DeLucia E.H. (2000) Xylem conductivity and vulnerability to cavitation of ponderosa pine growing in contrasting climates. Tree Physiology 20, 859–867.

Maherali H. & Pockman W.T. (2004) Adaptive Variation in the Vulnerability of Woody Plants To Xylem Cavitation. Ecology 85, 2184–2199.

Mantova M., Menezes-Silva P.E., Badel E., Cochard H. & Torres-Ruiz J.M. (2021) The interplay of hydraulic failure and cell vitality explains tree capacity to recover from drought. Physiologia Plantarum, ppl.13331.

Mardia K. V., Kent J.T. & Bibby J.M. (1979) Multivariate Analysis. Academic Press.

Martinez-Vilalta J., Anderegg W.R.L., Sapes G. & Sala A. (2019) Greater focus on water pools may improve our ability to understand and anticipate drought-induced mortality in plants. New Phytologist 223, 22–32.

Martinez-Vilalta J., Sala A. & Pinol J. (2004) The hydraulic architecture of Pinaceae. Plant Ecology 171, 3–13.

Marusig D., Petruzzellis F., Tomasella M., Napolitano R., Altobelli A. & Nardini A. (2020) Correlation of Field-Measured and Remotely Sensed Plant Water Status as a Tool to Monitor the Risk of Drought-Induced Forest Decline. Forests 11, 77.

Matías L., González-Díaz P. & Jump A.S. (2014) Larger investment in roots in southern range-edge populations of Scots pine is associated with increased growth and seedling resistance to extreme drought in response to simulated climate change. Environmental and Experimental Botany 105, 32–38.

Mcculloh K.A., Johnson D.M., Meinzer F.C. & Woodruff D.R. (2014) The dynamic pipeline: Hydraulic capacitance and xylem hydraulic safety in four tall conifer species. Plant, Cell and Environment 37, 1171–1183.

Meinzer F.C., Woodruff D.R., Marias D.E., Smith D.D., McCulloh K.A., Howard A.R. & Magedman A.L. (2016) Mapping ‘hydroscapes’ along the iso- to anisohydric continuum of stomatal regulation of plant water status. Ecology Letters.

Mencuccini M., Minunno F., Salmon Y., Martínez-Vilalta J. & Hölttä T. (2015) Coordination of physiological traits involved in drought-induced mortality of woody plants. New Phytologist 208, 396–409.

Mirzaie M., Darvishzadeh R., Shakiba A., Matkan A.A., Atzberger C. & Skidmore A. (2014) Comparative analysis of different uni- and multi-variate methods for estimation of vegetation water content using hyper-spectral measurements. International Journal of Applied Earth Observation and Geoinformation 26, 1–11.

Mischan M.M., Pinho S.Z. de & Carvalho L.R. de (2011) Determination of a point sufficiently close to the asymptote in nonlinear growth functions. Scientia Agricola 68, 109–114.

Mitchell P.J., O’Grady A.P., Tissue D.T., White D. a., Ottenschlaeger M.L. & Pinkard E. a. (2013) Drought response strategies define the relative contributions of hydraulic dysfunction and carbohydrate depletion during tree mortality. New Phytologist 197, 862–872.

Nardini A., Petruzzellis F., Marusig D., Tomasella M., Natale S., Altobelli A., … Zini L. (2020) Water ‘on the rocks’: a summer drink for thirsty trees? New Phytologist, 1–14.

O’Brien M.J., Leuzinger S., Philipson C.D., Tay J. & Hector A. (2014) Drought survival of tropical tree seedlings enhanced by non-structural carbohydrate levels. Nature Climate Change 4, 1–5.

Olson M.E., Soriano D., Rosell J.A., Anfodillo T., Donoghue M.J., Edwards E.J., … Méndez-Alonzo R. (2018) Plant height and hydraulic vulnerability to drought and cold. Proceedings of the National Academy of Sciences 115, 7551–7556.

Padilla F. & Pugnaire F. (2016) Rooting Depth and Soil Moisture Control Mediterranean Woody Seedling Survival during Drought. Functional Ecology 21, 489–495.

Piñol J. & Sala a. (2000) Ecological implications of xylem cavitation for several Pinaceae in the Pacific Northern USA. Functional Ecology 14, 538–545.

Potter K.M., Hipkins V.D., Mahalovich M.F. & Means R.E. (2013) Mitochondrial DNA haplotype distribution patterns in Pinus ponderosa (Pinaceae): Range-wide evolutionary history and implications for conservation. American Journal of Botany 100, 1562–1579.

Quentin A.G., Pinkard E.A., Ryan M.G., Tissue D.T., Baggett L.S., Adams H.D., … Woodruff D.R. (2015) Non-structural carbohydrates in woody plants compared among laboratories. Tree Physiology 35, 1146–1165.

R Core Team (2020) R: A language and environment for statistical computing.

Rao K., Anderegg W.R.L., Sala A., Martínez-Vilalta J. & Konings A.G. (2019) Satellite-based vegetation optical depth as an indicator of drought-driven tree mortality. Remote Sensing of Environment 227, 125–136.

Roskilly B., Keeling E., Hood S., Giuggiola A. & Sala A. (2019) Conflicting functional effects of xylem pit structure relate to the growth-longevity trade-off in a conifer species. Proceedings of the National Academy of Sciences 116, 15282–15287.

Saatchi S., Asefi-Najafabady S., Malhi Y., E. O. C. Aragão L., Anderson L.O., Myneni R.B. & Nemani R. (2013) Persistent effects of a severe drought on Amazonian forest canopy. Proceedings of the National Academy of Sciences 110, 565–570.

Sanders G.J. & Arndt S.K. (2012) Osmotic Adjustment Under Drought Conditions. In Plant Responses to Drought Stress. pp. 199–229. Springer Berlin Heidelberg, Berlin, Heidelberg.

Sapes G., Roskilly B., Dobrowski S., Maneta M., Anderegg W.R.L., Martinez-Vilalta J. & Sala A. (2019) Plant water content integrates hydraulics and carbon depletion to predict drought-induced seedling mortality. Tree Physiology 39, 1300–1312.

Sergent A.-S., Bréda N., Sanchez L., Bastein J.-C. & Rozenberg P. (2014) Coastal and interior Douglas-fir provenances differ in growth performance and response to drought episodes at adult age. Annals of Forest Science 71, 709–720.

Sevanto S., McDowell N.G., Dickman L.T., Pangle R. & Pockman W.T. (2014) How do trees die? A test of the hydraulic failure and carbon starvation hypotheses. Plant, cell & environment 37, 153–61.

Simeone C., Maneta M.P., Holden Z.A., Sapes G., Sala A. & Dobrowski S.Z. (2019) Coupled ecohydrology and plant hydraulics modeling predicts ponderosa pine seedling mortality and lower treeline in the US Northern Rocky Mountains. New Phytologist 221, 1814–1830.

Sperry J.S., Donnelly J.R. & Tyree M.T. (1988) A method for measuring hydraulic conductivity and embolism in xylem. Plant, Cell & Environment 11, 35–40.

Sperry J.S. & Love D.M. (2015) Tansley review What plant hydraulics can tell us about responses to climate-change droughts.

Stout D.L. & Sala A. (2003) Xylem vulnerability to cavitation in Pseudotsuga menziesii and Pinus ponderosa from contrasting habitats. Tree Physiology, 43–50.

Subbarao G.V., Nam N.H., Chauhan Y.S. & Johansen C. (2000) Osmotic adjustment, water relations and carbohydrate remobilization in pigeonpea under water deficits. Journal of Plant Physiology 157, 651–659.

Tognetti R., Michelozzi M. & Giovannelli a (1997) Geographical variation in water relations, hydraulic architecture and terpene composition of Aleppo pine seedlings from Italian provenances. Tree Physiology 17, 241–250.

Torres-Ruiz J.M., Jansen S., Choat B., McElrone A.J., Cochard H., Brodribb T.J., … Delzon S. (2015) Direct X-Ray Microtomography Observation Confirms the Induction of Embolism upon Xylem Cutting under Tension. Plant Physiology 167, 40–43.

Torres-Ruiz J.M., Sperry J.S. & Fernández J.E. (2012) Improving xylem hydraulic conductivity measurements by correcting the error caused by passive water uptake. Physiologia Plantarum 146, 129–135.

Trenberth K.E., Dai A., Van Der Schrier G., Jones P.D., Barichivich J., Briffa K.R. & Sheffield J. (2014) Global warming and changes in drought. Nature Climate Change 4, 17–22.

Trifilo P., Barbera P.M., Raimondo F., Nardini A. & Gullo M. a. L. (2014) Coping with drought-induced xylem cavitation: coordination of embolism repair and ionic effects in three Mediterranean evergreens. Tree Physiology 34, 109–122.

Tyree M.T., Engelbrecht B.M.J., Vargas G., Kursar T. a, States U., Forest A., … Vermont M.T.T. (2003) Desiccation Tolerance of Five Tropical Seedlings in Panama. Relationship to a Field Assessment of Drought Performance. Plant physiology 132, 1439–1447.

Tyree M.T. & Sperry J.S. (1989) Vulnerability of Xylem to Cavitation and Embolism. Ann. Rev. Plant. Phys. Mol. Bio. 40, 19–38.

Tyree M.T., Vargas G., Engelbrecht B.M.J. & Kursar T. a (2002) Drought until death do us part: a case study of the desiccation-tolerance of a tropical moist forest seedling-tree, Licania platypus (Hemsl.) Fritsch. Journal of experimental botany 53, 2239–2247.

Ullah S., Skidmore A.K., Naeem M. & Schlerf M. (2012) An accurate retrieval of leaf water content from mid to thermal infrared spectra using continuous wavelet analysis. Science of the Total Environment 437, 145–152.

Umaña M. & Swenson N. (2019) Does trait variation within broadly distributed species mirror patterns across species? A case study in Puerto Rico. 100, 1–11.

Walker S.H. & Duncan D.B. (1967) Estimation of the probability of an event as a function of several independent variables. Biometrika.

Wang Q. & Li P. (2012) Identification of robust hyperspectral indices on forest leaf water content using PROSPECT simulated dataset and field reflectance measurements. Hydrological Processes 26, 1230–1241.

Zhang Y., Zhou S., Gentine P. & Xiao X. (2019) Can vegetation optical depth reflect changes in leaf water potential during soil moisture dry-down events? Remote Sensing of Environment 234, 1–10.

