## Supplementary material for "Relative water content consistently predicts drought mortality risk in seedling populations with different morphology, physiology, and times to death": Fig. S1

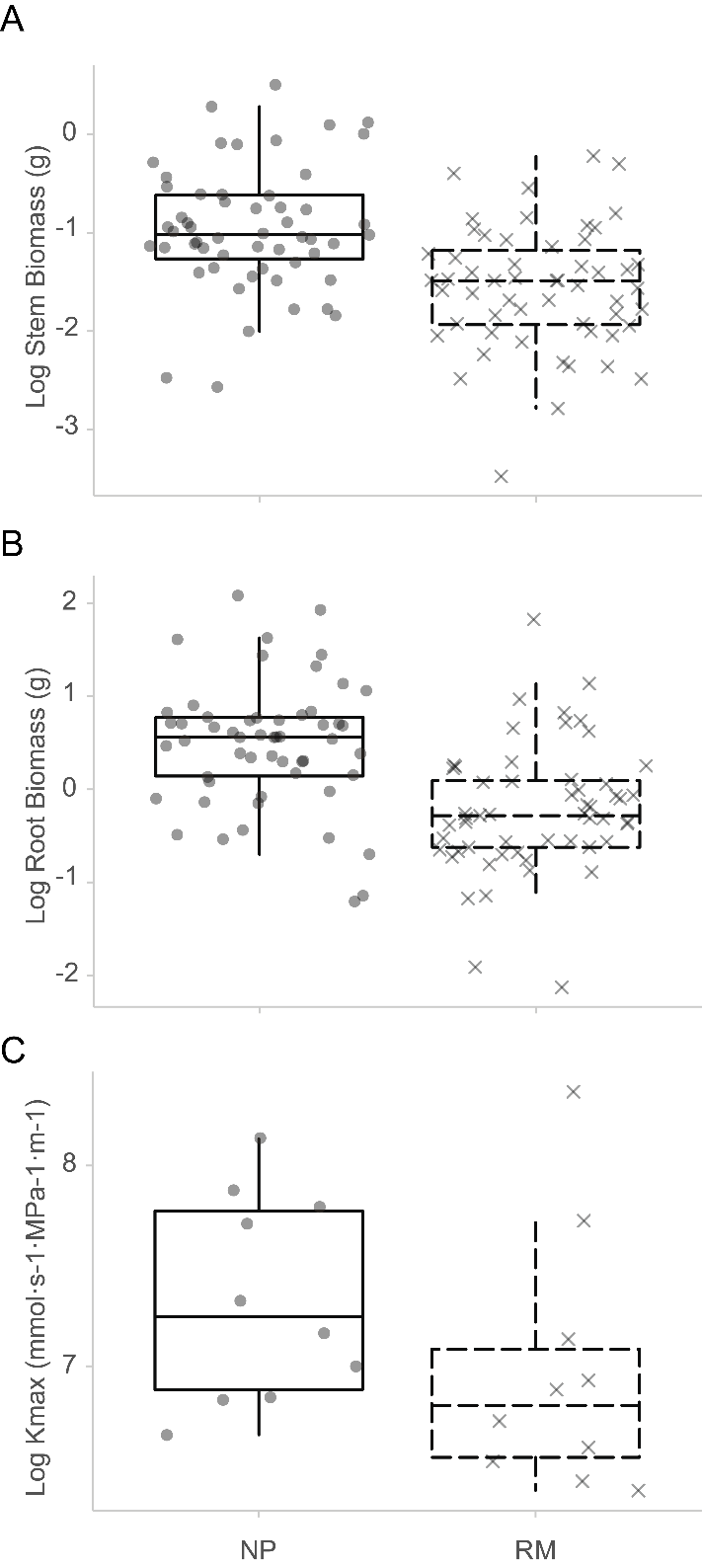

**Figure S1.** Differences in stem biomass, root biomass, and maximum stem hydraulic conductivity between North Plateau (NP) and Rocky Mountain (RM) seedlings. Differences among populations are significant across all panels.
