## Supplementary material for "Relative water content consistently predicts drought mortality risk in seedling populations with different morphology, physiology, and times to death": Fig. S2

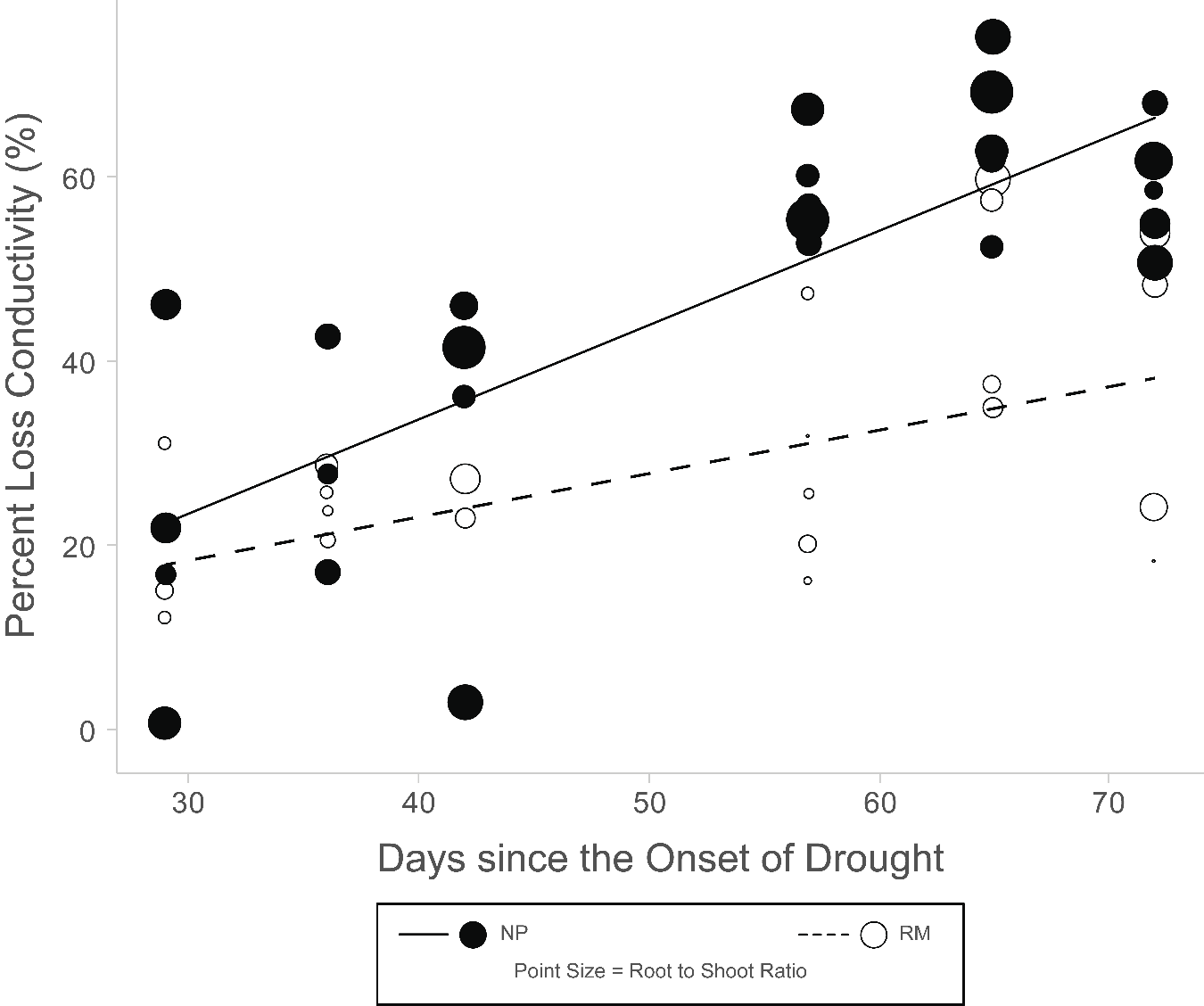

**Figure S2.** Differences in wood percent loss of hydraulic conductivity (PLC) over time between populations were associated with differences in biomass allocation. Plants with greater root to shoot ratio experienced PLC at faster rates. North Plateau (NP) and Rocky Mountain (RM) seedlings are shown by solid circles and solid lines, and by open circles and dashed lines, respectively. The size of the points represents the root to shoot ratio value.
