## Supplementary material for "Relative water content consistently predicts drought mortality risk in seedling populations with different morphology, physiology, and times to death": Methods S1

**Methods S1.** Description of methods used for morphological measurements and non-structural carbohydrates (NSC).

**Morphological measurements**

Upon harvest, we measured plant height and root system length in each seedling as the distance from the root collar to the highest needle and to the end of the longest root, respectively. We calculated plant length as the sum of plant height and root system length. We took pictures of the canopy of each seedling and estimate total canopy leaf area using ImageJ software 1.48 (National Institutes of Health). Organ dry weights were used to calculate whole plant biomass as the sum of needle, stem, and root dry weights. Root to shoot ratios were calculated as root dry weight divided by the sum of leaf and stem dry weights.

**Non-structural carbohydrates**

Sample powder was dissolved in 1.6 mL of distilled water and incubated at 100 °C for 60 min to extract carbohydrates. An aliquot of the extract was used to determine soluble sugar concentrations (i.e., glucose, fructose, and sucrose) through enzymatic conversion of sucrose and fructose into glucose by invertase from *Saccharomyces cerevisiae* and phosphoglucose isomerase, respectively (I4504 and G3293, Sigma-Aldrich). Total *NSC* concentration was obtained from another aliquot incubated in amyloglucosidase from *Aspergillus niger* (10115, Sigma-Aldrich) at 50 °C during 16 hours to break down all *NSC* (starch included) into glucose. In both cases, the concentration of glucose was determined photometrically in a 96-well microplate reader (BioTek™ EL800, Winooski, United States) after enzymatic conversion of glucose into glucose-6-phosphate by glucose hexokinase. The dehydrogenation of glucose causes an increase in optical density at 340 nm. All the NSC and soluble sugar concentrations were expressed as percentages of dry matter. Although the quantification of NSC has been proven difficult and inconsistent among laboratories, the reasonable consistency within a given laboratory allows comparisons among samples (Quentin *et al.* 2015). We calculated the total pool of NSCs, starch, soluble sugars, and glucose or fructose in each tissue by multiplying the corresponding concentration of each tissue by its dry weight. Concentrations (total NSC and each individual component) were later scaled up to whole-plant level as explained in the RWC section. Finally, the relative degree of NSC depletion of each individual (here on percent NSC deviation from control) was calculated at both tissue and whole-plant levels by subtracting the mean NSC concentration of LL seedlings (i.e. controls) from each individual value and dividing that difference by the mean. This ratio was then multiplied by 100 to represent it as a percentage. Percent NSC deviation from control can range from negative to positive values with negative values expressing NSC depletion and positive values expressing greater NSC pools than the average non-stressed seedling (Adams *et al.* 2017).

Adams H.D., Zeppel M.J.B., Anderegg W.R.L., Hartmann H., Landhäusser S.M., Tissue D.T., … McDowell N.G. (2017) A multi-species synthesis of physiological mechanisms in drought-induced tree mortality. *Nature Ecology & Evolution* **1**, 1285–1291.

Quentin A.G., Pinkard E.A., Ryan M.G., Tissue D.T., Baggett L.S., Adams H.D., … Woodruff D.R. (2015) Non-structural carbohydrates in woody plants compared among laboratories. *Tree Physiology* **35**, 1146–1165.
