## Supplementary material for "Relative water content consistently predicts drought mortality risk in seedling populations with different morphology, physiology, and times to death": Methods S2

**Methods S2.** Derivative explanation and steps to obtain a threshold consistency index.

First, we built a generalized linear model (GLM) with a binomial distribution that estimates the probability of mortality as a function of a predictor. Then, we extract the equation describing the fit line of the GLM model (teal) and use it to calculate the 1^st^ (grey), 2^nd^ (gold), and 3^rd^ (red) derivatives of that function.

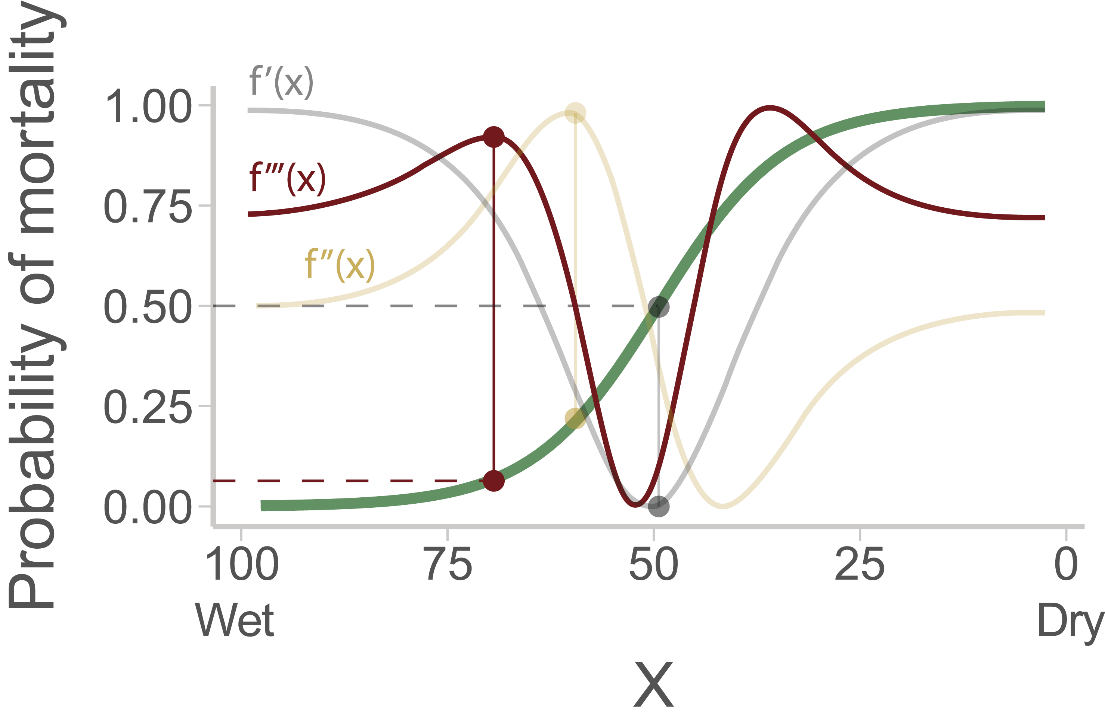

The extrema (maxima and minima) in these derivatives can be used to identify key points in the mortality curve.

The 1^st^ derivative (grey) describes changes in the rate at which mortality increases as physiological status declines, equivalent to a “velocity” of mortality risk. The minimum of the first derivative (grey dot) identifies the inflection point (grey vertical line) on the mortality function (green), when mortality is increasing at its maximum rate. This coincides with the value at 50% mortality risk (dashed grey line).

The 2^nd^ derivative (gold) describes how the velocity of mortality risk changes as physiological status declines, equivalent to the “acceleration” of mortality risk. The maxima (gold dot) and minima of the 2^nd^ derivative identify the points in the mortality curve where the acceleration of mortality is greatest or lowest, respectively, which delimit the middle section of the non-linear portion of the mortality function. Hence, the 2^nd^ derivative does not provide the point at which mortality starts to accelerate.

The 3^rd^ derivative (red) describes how the acceleration of mortality risk changes as physiological status declines. Maxima and minima of the 3^rd^ derivative identify the points in the mortality curve where the change in acceleration is most suddenly increasing or decreasing. A maximum or minimum peak (depending on the nature of the physiological predictor) near the wet end of the range of physiological predictor values (X axis) identifies the point on the mortality curve at which there is a sudden increase in mortality risk (red dot). This is the incipient mortality threshold. We can assess the degree of incipiency of this threshold (how well it captures the early stages of mortality) by projecting the incipient threshold onto the Y axis (dashed red line) and evaluating how close the probability of mortality at the threshold is from 0. When the probability of mortality at the physiological predictor threshold is close to zero it indicates that the predictor captures the very early stages of mortality (incipient mortality). In contrast, if the probability of mortality at the threshold is for instance 20%, it indicates that the physiological predictor is not capturing the early stages of mortality.

Estimating threshold consistency across organs and races.

The consistency index assesses mortality thresholds in terms of their X axis value, the value of the physiological predictor (i.e. how consistent the physiological predictor is). We obtain a consistency index through the following steps. For a given predictor, we first calculate the average value of the predictor at the mortality threshold across all mortality curves. Since we have mortality curves for each organ within a population, the average threshold value is calculated as:

$$\bar{Threshold}= \frac{\begin{aligned} {(Threshold}_{Leaf NP}+ {Threshold}_{Stem NP}+ {Threshold}_{Root NP}+ {Threshold}_{Plant NP}+ \\ {Threshold}_{Leaf RM}+ {Threshold}_{Stem RM}+ {Threshold}_{Root RM}+ {Threshold}_{Plant RM}) \end{aligned}}{Total number of thresholds}$$

Where NP and RM are the North Plateau and Rocky Mountain populations, respectively. Second, we calculate how each threshold used in the previous equation deviates from the average threshold (absolute value of that deviation). For instance, to calculate how the threshold corresponding to the whole plant mortality curve within the NP population deviates from the average threshold value, we do the following calculation:

$${Threshold deviation}_{Plant NP}= \left| {Threshold}_{Plant NP}- \bar{Threshold} \right|$$

Third, to make values comparable across physiological predictors with different units, we divide the absolute deviation by the average threshold value to obtain a standard, unitless index.

$${Standardized threshold deviation}_{Plant NP}= \frac{{Threshold deviation}_{Plant NP}}{\bar{Threshold}}$$

Because this calculation is a ratio, values can become disproportionately high when the denominator (mean threshold value) is closer to 0. To reduce this effect, we add a logarithmic transformation to normalize the distributions of values among predictors and make them more comparable. In this index, less negative values indicate more variable thresholds. Last, to make it a measure of consistency (greater values corresponding to greater consistency) rather than variability we multiply this index by -1.

$$Threshold consistency index=-1* log({Standardized threshold variability}_{Plant NP})$$
