## Supplementary material for "Relative water content consistently predicts drought mortality risk in seedling populations with different morphology, physiology, and times to death": Methods S3

**Methods S3.** Relative water content (RWC) mortality curves for each organ (green = leaves; yellow = stem; black = root; purple = whole plant) within the North Plateau race. For each organ, the 1^st^ (black), 2^nd^ (gold), and 3^rd^ (red) derivatives are shown. Vertical black, gold, and red lines indicate the inflection point, the point of maximum acceleration, and the incipient mortality threshold of the mortality curve, respectively. Dashed horizontal lines indicate the mortality value at the threshold (red) and 5% mortality (blue) as a visual reference. The farther apart the red and blue horizontal dashed lines are, the less incipient the threshold is. In this case the two lines overlap to produce a dark color.

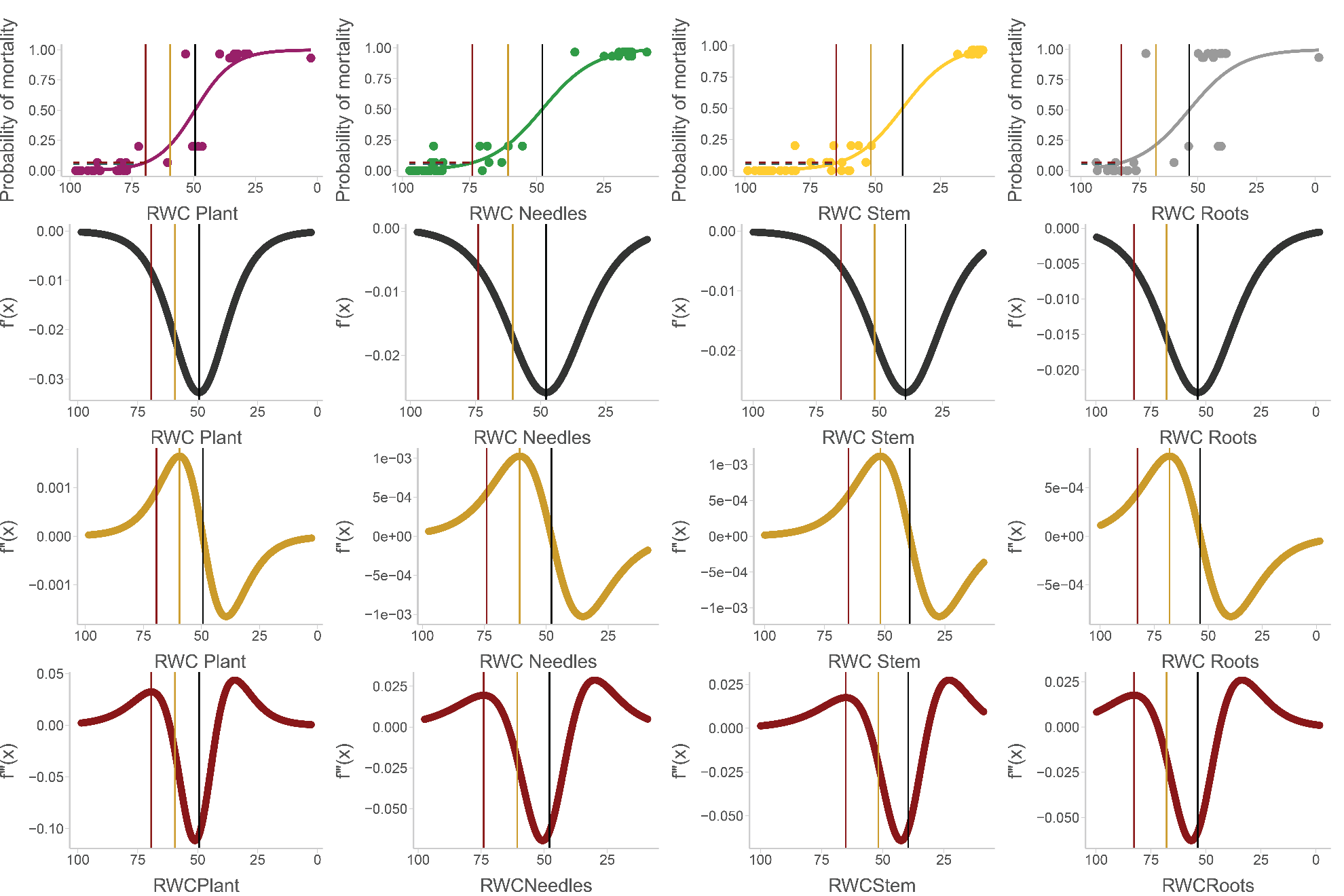

Relative water content (RWC) mortality curves for each organ (green = leaves; yellow = stem; black = root; purple = whole plant) within the Rocky Mountain race. For each organ, the 1^st^ (black), 2^nd^ (gold), and 3^rd^ (red) derivatives are shown. Vertical black, gold, and red lines indicate the inflection point, the point of maximum acceleration, and the incipient mortality threshold of the mortality curve, respectively. Dashed horizontal lines indicate the mortality value at the threshold (red) and 5% mortality (blue) as a visual reference. The farther apart the red and blue horizontal dashed lines are, the less incipient the threshold is. In this case the two lines overlap to produce a dark color.

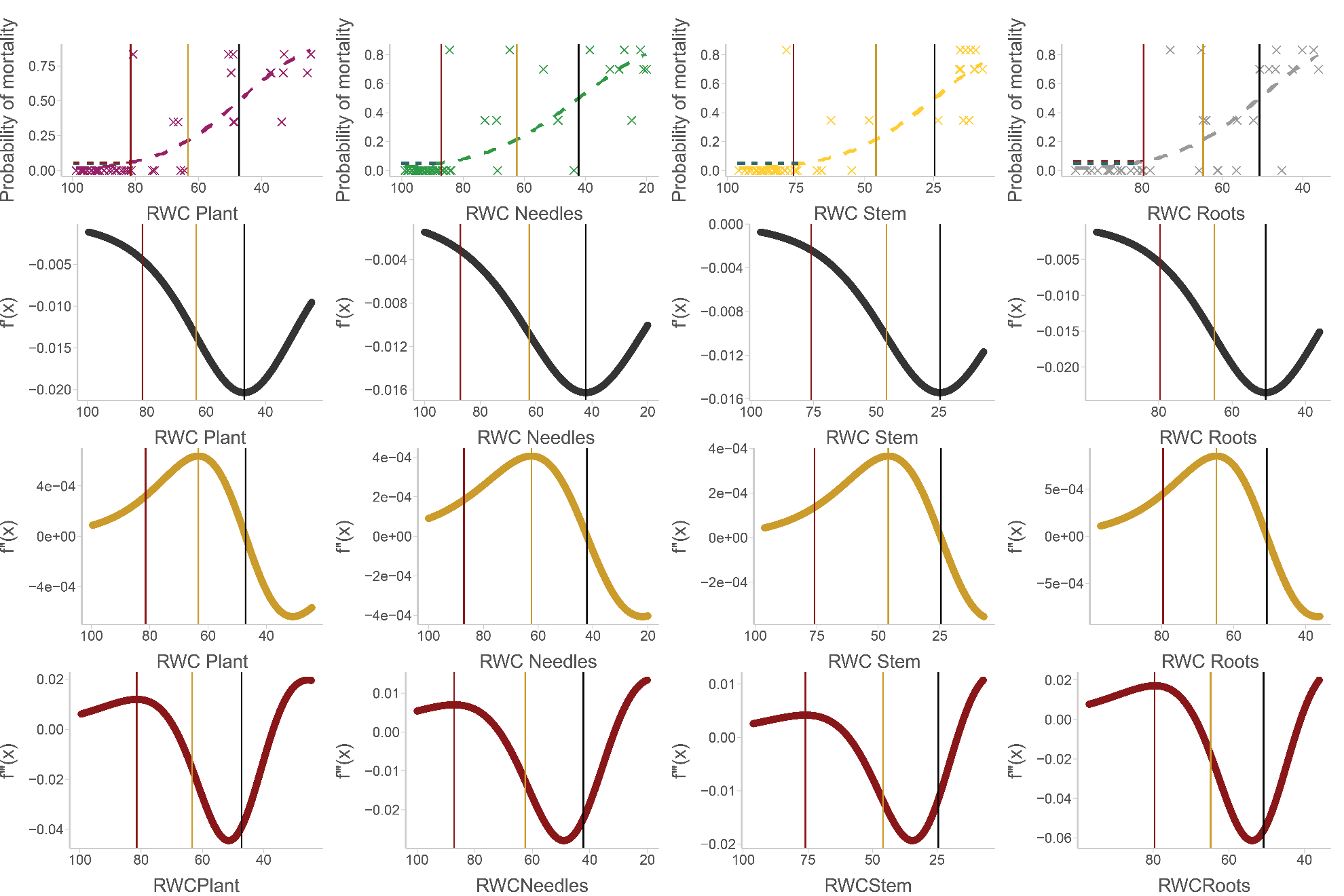

Water potential (WP) mortality curves for each organ (green = leaves; yellow = stem) within the North Plateau race. For each organ, the 1^st^ (black), 2^nd^ (gold), and 3^rd^ (red) derivatives are shown. Vertical black, gold, and red lines indicate the inflection point, the point of maximum acceleration, and the incipient mortality threshold of the mortality curve, respectively. Dashed horizontal lines indicate the mortality value at the threshold (red) and 5% mortality (blue) as a visual reference. The farther apart the red and blue horizontal dashed lines are, the less incipient the threshold is.

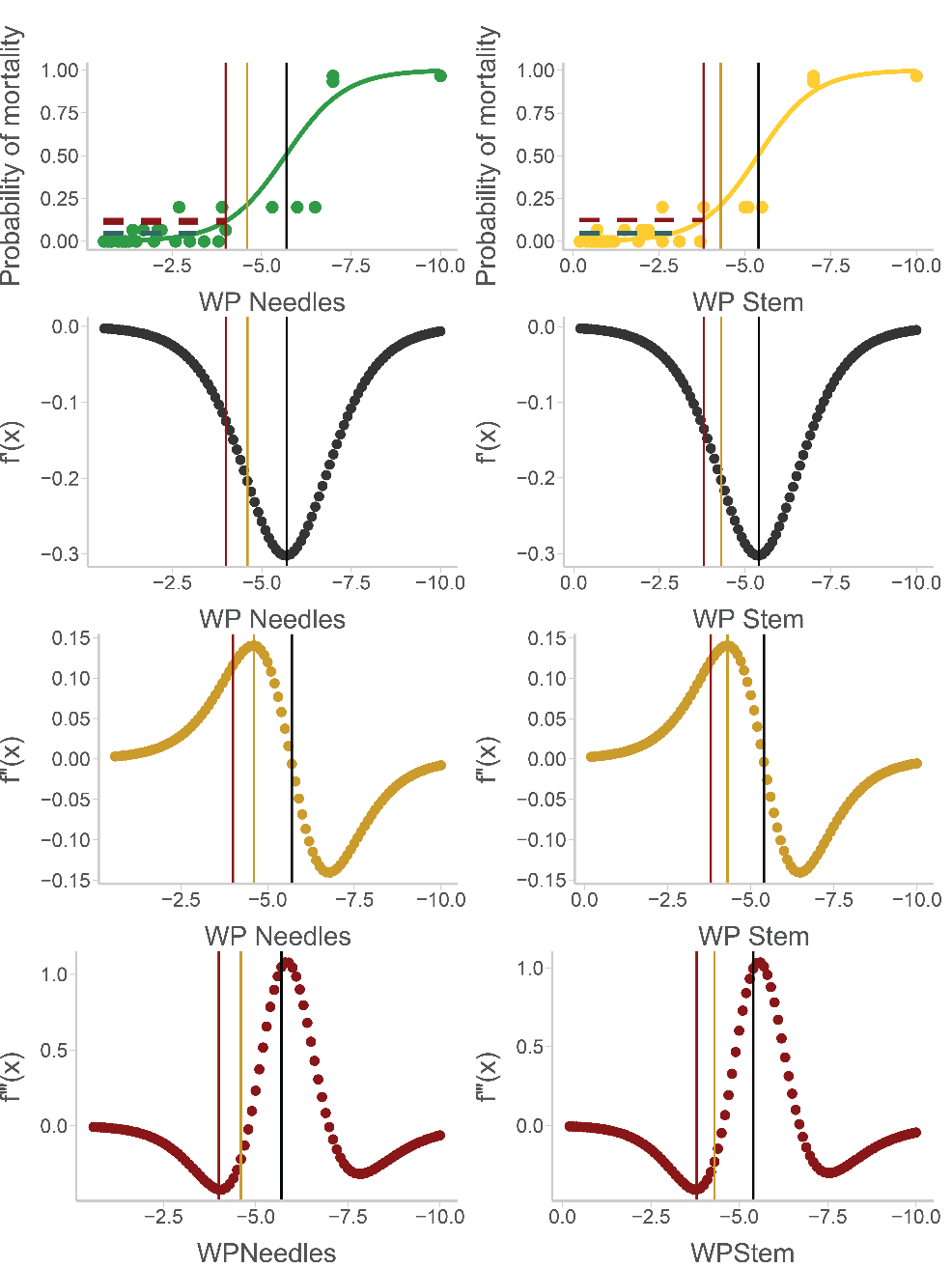

Water potential (WP) mortality curves for each organ (green = leaves; yellow = stem) within the Rocky Mountain race. 1^st^ (black), 2^nd^ (gold), and 3^rd^ (red) derivatives. For each organ, the 1^st^ (black), 2^nd^ (gold), and 3^rd^ (red) derivatives are shown. Vertical black, gold, and red lines indicate the inflection point, the point of maximum acceleration, and the incipient mortality threshold of the mortality curve, respectively. Dashed horizontal lines indicate the mortality value at the threshold (red) and 5% mortality (blue) as a visual reference. The farther apart the red and blue horizontal dashed lines are, the less incipient the threshold is.

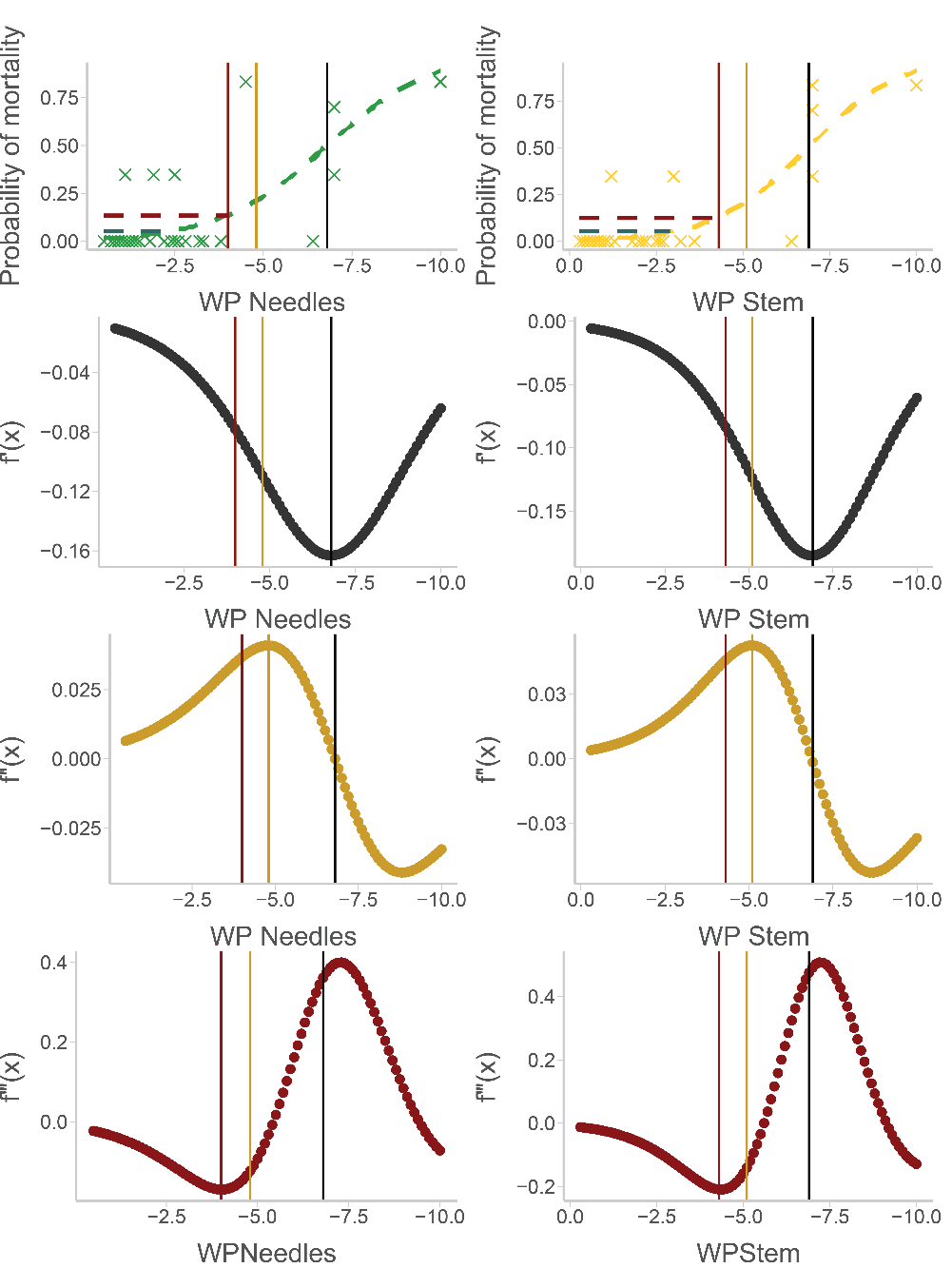
