## Supplementary material for "Relative water content consistently predicts drought mortality risk in seedling populations with different morphology, physiology, and times to death": Table S1

**Table S1.** Number of percent loss of conductivity (PLC) data points removed by North Plateau (NP) and Rocky Mountain (RM) populations, organ and sampling group. The number of points removed was similar in each population and organ. The highest number of points removed in each organ corresponds to Controls. Removed data within this group correspond to two groups of well-watered control seedlings (preconditioned and non-preconditioned) that were measured at the end of the experiment. These seedlings grew over time which resulted in greater hydraulic conductivity than in the preconditioned controls used to calculate population-level Kmax at day 0. Removal of these data is unlikely to drive patterns found in PLC between populations given that observed differences between populations appeared at late stages of drought.

| **Group** | **Stem** | | | **Root** | | | **Wood** | | |
| --- | --- | --- | --- | --- | --- | --- | --- | --- | --- |
|  | **NP** | **RM** | **Difference** | **NP** | **RM** | **Difference** | **NP** | **RM** | **Difference** |
| Controls | 9 | 5 | 4 | 6 | 3 | 3 | 9 | 4 | 5 |
| Drought - Day 29 | 2 | 5 | -3 | 1 | 2 | -1 | 1 | 2 | -1 |
| Drought - Day 36 | 2 | 1 | 1 | 2 | 1 | 1 | 2 | 1 | 1 |
| Drought - Day 42 | 2 | 3 | -1 | 1 | 3 | -2 | 1 | 3 | -2 |
| Drought - Day 57 | 0 | 0 | 0 | 0 | 0 | 0 | 0 | 0 | 0 |
| Drought - Day 65 | 0 | 1 | -1 | 0 | 0 | 0 | 0 | 1 | -1 |
| Drought - Day 72 | 0 | 0 | 0 | 0 | 0 | 0 | 0 | 0 | 0 |
| **TOTAL** | **15** | **15** |  | **10** | **9** |  | **13** | **11** |  |
