## Supplementary material for "Relative water content consistently predicts drought mortality risk in seedling populations with different morphology, physiology, and times to death": Table S2

**Table S2.** Linear models assessing the effect of plant length on hydraulic conductivity and conductance. Response variables were logarithmically transformed to meet model assumptions. Population categories correspond to North Plateau (NP) and Rocky mountain (RM).

| **Model and Factors** | **Estimate** | **95% C.I. Estimates** | | ***p-value*** | **d.f. (res.)** | **Adjusted R square** |
| --- | --- | --- | --- | --- | --- | --- |
|  |  | **2.5%** | **97.5%** |  |  |  |
| *Log(Stem Hydraulic Conductivity) = Plant length x Population* |  |  |  | **<0.001** | 102 | 0.18 |
| Intercept | 1.830401 | 1.37624098 | 2.28456091 | **<0.001** | - | - |
| *Plant Length* | 0.017699 | 0.01068521 | 0.02471359 | **<0.001** | - | - |
| *Population - RM* | 0.162318 | -0.01802708 | 0.34266243 | 0.077 | - | - |
| *Log(Root Hydraulic Conductance) = Plant length x Population* |  |  |  | **<0.001** | 105 | 0.08 |
| Intercept | 3.626437 | 3.265981513 | 3.98689211 | **<0.001** | - | - |
| *Plant Length* | 0.008438 | 0.002852591 | 0.01402426 | **0.003** | - | - |
| *Population - RM* | 0.148013 | 0.001958142 | 0.29406819 | **0.047** | - | - |
