## Supplementary material for "Relative water content consistently predicts drought mortality risk in seedling populations with different morphology, physiology, and times to death": Table S3

**Table S3.** Models assessing changes in drought intensity, hydraulic conductivity, NSC changes, degree of dehydration, canopy activity levels and population-level mortality over time starting at day 29 since the onset of drought. Population categories correspond to North Plateau (NP) and Rocky mountain (RM). Only significant factors are shown. A logistic model was used for probability of mortality because it could not be transformed to meet linear model assumptions. Probability of mortality was also transformed to per unit basis following requirements of models with binomial distributions.

| **Model and Factors** | **Model type** | **Estimate** | **95% C.I. Estimates** | | ***p-value*** | **d.f. (res.)** | **Adjusted R square** |
| --- | --- | --- | --- | --- | --- | --- | --- |
|  |  |  | **2.5%** | **97.5%** |  |  |  |
| *Soil Water Potential = Days since the 29th day of Drought x Population* | LM |  |  |  | **<0.001** | 57 | 0.30 |
| Intercept |  | -1.075160 | -0.239196239 | 0.48603325 | **0.031** | - | - |
| *Days since the 29th day of Drought* |  | -0.041689 | -0.073496211 | -0.05567413 | **<0.001** | - | - |
| *Population-RM* |  | 0.620238 | -0.503905295 | 0.52172409 | **0.029** | - | - |
| *Leaf Water Potential = Days since the 29th day of Drought x Population* | LM |  |  |  | **<0.001** | 53 | 0.77 |
| Intercept |  | 2.22232 | 0.95229030 | 3.4923595 | **<0.001** | - | - |
| *Days since the 29th day of Drought* |  | -0.15796 | -0.18092399 | -0.1350017 | **<0.001** | - | - |
| *Population-RM* |  | 0.79490 | 0.07352279 | 1.5162857 | **0.031** | - | - |
| *Stem Water Potential = Days since the 29th day of Drought x Population* | LM |  |  |  | **<0.001** | 54 | 0.83 |
| Intercept |  | 3.15997 | 2.04923568 | 4.2706960 | **<0.001** | - | - |
| *Days since the 29th day of Drought* |  | -0.17223 | -0.19250203 | -0.1519484 | **<0.001** | - | - |
| *Population-RM* |  | 0.59747 | -0.03882642 | 1.2337579 | 0.065 | - | - |

| **Model and Factors** | **Model type** | **Estimate** | **95% C.I. Estimates** | | ***p-value*** | **d.f. (res.)** | **Adjusted R square** |
| --- | --- | --- | --- | --- | --- | --- | --- |
|  |  |  | **2.5%** | **97.5%** |  |  |  |
| *Wood PLC = Days since the 29th day of Drought x Population* | LM |  |  |  | **<0.001** | 44 | 0.57 |
| Intercept |  | -7.5348 | -25.9228774 | 10.87328814 | 0.414 | - | - |
| *Days since the 29th day of Drought* |  | 1.0279 | 0.6910973 | 1.36463821 | **<0.001** | - | - |
| *Population-RM* |  | 11.9945 | -14.0948133 | 38.08376569 | 0.360 | - | - |
| *Days since the 29th day of Drought x Population-RM* |  | -0.5622 | -1.0405618 | -0.08380609 | **0.022** | - | - |
| *Wood PLC = Days since the 29th day of Drought x Population x Root Shoot Ratio* | LM |  |  |  | **<0.001** | 43 | 0.60 |
| Intercept |  | -17.6897 | -37.7473072 | 2.36782551 | 0.082 | - | - |
| *Days since the 29th day of Drought* |  | 0.9740 | 0.6460260 | 1.30191448 | **<0.001** | - | - |
| *Population-RM* |  | 17.6961 | -7.9630927 | 43.35522894 | 0.172 | - | - |
| *Root Shoot Ratio* |  | 8.3719 | 0.6118301 | 16.13193591 | **0.035** | - | - |
| *Days since the 29th day of Drought x Population-RM* |  | -0.5591 | -1.0194898 | -0.09866543 | **0.018** | - | - |
| *Stem PLC = Days since the 29th day of Drought x Population* | LM |  |  |  | **<0.001** | 40 | 0.49 |
| Intercept |  | 8.1380 | -15.6972775 | 31.973188 | 0.495 | - | - |
| *Days since the 29th day of Drought* |  | 1.2206 | 0.7976874 | 1.643427 | **<0.001** | - | - |
| *Population-RM* |  | -24.8821 | -37.5015720 | -12.262635 | **<0.001** | - | - |
| *Root PLC = Days since the 29th day of Drought x Population* | LM |  |  |  | **<0.001** | 45 | 0.27 |
| Intercept |  | 9.1774 | -17.7046217 | 36.059357 | 0.496 | - | - |
| *Days since the 29th day of Drought* |  | 1.0530 | 0.5609697 | 1.545101 | **<0.001** | - | - |
| *Population-RM* |  | 39.2267 | 1.1932278 | 77.260242 | **0.044** | - | - |
| *Days since the 29th day of Drought x Population-RM* |  | -0.9189 | -1.6133898 | -0.224493 | **0.011** | - | - |

| **Model and Factors** | **Model type** | **Estimate** | **95% C.I. Estimates** | | ***p-value*** | **d.f. (res.)** | **Adjusted R square** |
| --- | --- | --- | --- | --- | --- | --- | --- |
|  |  |  | **2.5%** | **97.5%** |  |  |  |
| *Plant NSC Relative to Control= Days since the 29th day of Drought x Population* | LM |  |  |  | **<0.001** | 57 | 0.57 |
| Intercept |  | 16.0352 | 1.951829 | 30.1186031 | **0.026** | - | - |
| *Days since the 29th day of Drought* |  | -1.0015 | -1.258506 | -0.7445081 | **<0.001** | - | - |
| *Population-RM* |  | -17.7042 | -25.718929 | -9.6894772 | **<0.001** | - | - |
| *Leaf NSC Relative to Control = Days since the 29th day of Drought x Population* | LM |  |  |  | **<0.001** | 58 | 0.16 |
| Intercept |  | -9.8766 | -29.872194 | 10.1190894 | **0.327** | - | - |
| *Days since the 29th day of Drought* |  | -0.6663 | -1.046874 | -0.2856296 | **<0.001** | - | - |
| *Stem NSC Relative to Control = Days since the 29th day of Drought x Population* | LM |  |  |  | **0.001** | 57 | 0.38 |
| Intercept |  | 41.9929 | 17.214449 | 66.7712653 | **<0.001** | - | - |
| *Days since the 29th day of Drought* |  | -0.9816 | -1.433736 | -0.5294045 | **<0.001** | - | - |
| *Population-RM* |  | -30.3919 | -44.493048 | -16.2907208 | **<0.001** |  |  |
| *Root NSC Relative to Control = Days since the 29th day of Drought x Population* | LM |  |  |  | **<0.001** | 56 | 0.70 |
| Intercept |  | 59.8135 | 37.2986770 | 82.328291 | **<0.001** | - | - |
| *Days since the 29th day of Drought* |  | -1.6364 | -2.0650049 | -1.207855 | **<0.001** | - | - |
| *Population-RM* |  | -75.7772 | -107.6179209 | -43.936431 | **<0.001** | - | - |
| *Days since the 29th day of Drought x Population-RM* |  | 0.7657 | 0.1595556 | 1.371749 | **0.014** | - | - |

| **Model and Factors** | **Model type** | **Estimate** | **95% C.I. Estimates** | | ***p-value*** | **d.f. (res.)** | **Adjusted R square** |
| --- | --- | --- | --- | --- | --- | --- | --- |
|  |  |  | **2.5%** | **97.5%** |  |  |  |
| *Plant Starch = Days since the 29th day of Drought x Population* | LM |  |  |  | **0.007** | 52 | 0.11 |
| Intercept |  | 0.5500 | 0.2304901 | 0.8694543 | **0.001** | - | - |
| *Population-RM* |  | 0.6194 | 0.1756866 | 1.0630355 | **0.007** | - | - |
| *Leaf Starch = Days since the 29th day of Drought x Population* | LM |  |  |  | 0.068 | 53 | 0.04 |
| Intercept |  | 0.8127 | 0.36965578 | 1.255810 | **<0.001** | - | - |
| *Population-RM* |  | 0.5762 | -0.04474468 | 1.197226 | 0.068 | - | - |
| *Stem Starch = Days since the 29th day of Drought x Population* | LM |  |  |  | **0.003** | 56 | 0.16 |
| Intercept |  | 1.109684 | 0.44783451 | 1.771534086 | **0.001** | - | - |
| *Days since the 29th day of Drought* |  | -0.015056 | -0.02698653 | -0.003125525 | **0.014** | - | - |
| *Population-RM* |  | 0.466922 | 0.09757230 | 0.836271977 | **0.014** | - | - |
| *Root Starch = Days since the 29th day of Drought x Population* | LM |  |  |  | **0.004** | 56 | 0.17 |
| Intercept |  | 0.208537 | -0.90809186 | 1.3251654408 | 0.710 | - | - |
| *Days since the 29th day of Drought* |  | 0.003867 | -0.01738783 | 0.0251227766 | 0.717 | - | - |
| *Population-RM* |  | 2.174318 | 0.59516663 | 3.7534693930 | **0.007** | - | - |
| *Days since the 29th day of Drought x Population-RM* |  | -0.029074 | -0.05913355 | 0.0009855313 | 0.058 | - | - |

| **Model and Factors** | **Model type** | **Estimate** | **95% C.I. Estimates** | | ***p-value*** | **d.f. (res.)** | **Adjusted R square** |
| --- | --- | --- | --- | --- | --- | --- | --- |
|  |  |  | **2.5%** | **97.5%** |  |  |  |
| *Plant Sucrose = Days since the 29th day of Drought x Population* | LM |  |  |  | **<0.001** | 55 | 0.49 |
| Intercept |  | 3.276542 | 2.7440109 | 3.80907389 | **<0.001** | - | - |
| *Days since the 29th day of Drought* |  | -0.038013 | -0.0483597 | -0.02766645 | **<0.001** | - | - |
| *Leaf Sucrose = Days since the 29th day of Drought x Population* | LM |  |  |  | **<0.001** | 55 | 0.25 |
| Intercept |  | 3.03618 | 2.19551465 | 3.87685298 | **<0.001** | - | - |
| *Days since the 29th day of Drought* |  | -0.03605 | -0.05238754 | -0.01972061 | **<0.001** | - | - |
| *Stem Sucrose = Days since the 29th day of Drought x Population* | LM |  |  |  | **<0.001** | 58 | 0.18 |
| Intercept |  | 4.04918 | 2.96609793 | 5.13225850 | **<0.001** | - | - |
| *Days since the 29th day of Drought* |  | -0.03903 | -0.05964643 | -0.01841302 | **<0.001** | - | - |
| *Root Sucrose = Days since the 29th day of Drought x Population* | LM |  |  |  | **<0.001** | 57 | 0.37 |
| Intercept |  | 2.82421 | 2.08461089 | 3.56380943 | **<0.001** | - | - |
| *Days since the 29th day of Drought* |  | -0.03479 | -0.04829132 | -0.02129834 | **<0.001** | - | - |
| *Population-RM* |  | 0.64802 | 0.22711992 | 1.06891817 | **0.003** | - | - |

| **Model and Factors** | **Model type** | **Estimate** | **95% C.I. Estimates** | | ***p-value*** | **d.f. (res.)** | **Adjusted R square** |
| --- | --- | --- | --- | --- | --- | --- | --- |
|  |  |  | **2.5%** | **97.5%** |  |  |  |
| *Plant Glucose+Fructose = Days since the 29th day of Drought x Population* | LM |  |  |  | **0.050** | 57 | 0.05 |
| Intercept |  | 2.471464 | 1.84993843 | 3.09299 | **<0.001** | - | - |
| *Days since the 29th day of Drought* |  | -0.011756 | -0.02353408 | 2.20714e-05 | **0.050** | - | - |
| *Leaf Glucose+Fructose = Days since the 29th day of Drought x Population* | LM |  |  |  | **<0.001** | 58 | 0.31 |
| Intercept |  | 3.3822 | 2.949800 | 3.8145409 | **<0.001** | - | - |
| *Population-RM* |  | -1.5921 | -2.203517 | -0.9805892 | **<0.001** | - | - |
| *Stem Glucose+Fructose = Days since the 29th day of Drought x Population* | LM |  |  |  | 0.598 | 57 | 0.01 |
| Intercept |  | 2.026668 | 1.19452413 | 2.85881265 | **<0.001** | - | - |
| *Days since the 29th day of Drought* |  | -0.004173 | -0.01994241 | 0.01159629 | 0.598 | - | - |
| *Root Glucose+Fructose = Days since the 29th day of Drought x Population* | LM |  |  |  | **<0.001** | 58 | 0.24 |
| Intercept |  | 3.138640 | 2.40577928 | 3.87150124 | **<0.001** | - | - |
| *Days since the 29th day of Drought* |  | -0.030697 | -0.04464692 | -0.01674654 | **<0.001** | - | - |

| **Model and Factors** | **Model type** | **Estimate** | **95% C.I. Estimates** | | ***p-value*** | **d.f. (res.)** | **Adjusted R square** |
| --- | --- | --- | --- | --- | --- | --- | --- |
|  |  |  | **2.5%** | **97.5%** |  |  |  |
| *Plant RWC = Days since the 29th day of Drought x Population* | LM |  |  |  | **<0.001** | 54 | 0.71 |
| Intercept |  | 110.9513 | 99.456181 | 122.4465168 | **<0.001** | - | - |
| *Days since the 29th day of Drought* |  | -1.1934 | -1.403436 | -0.9833056 | **<0.001** | - | - |
| *Population-RM* |  | 12.6950 | 6.098499 | 19.2915497 | **<0.001** | - | - |
| *Leaf RWC = Days since the 29th day of Drought x Population* | LM |  |  |  | **<0.001** | 54 | 0.78 |
| Intercept |  | 141.8941 | 124.20776335 | 159.580410 | **<0.001** | - | - |
| *Days since the 29th day of Drought* |  | -1.8720 | -2.20868295 | -1.535356 | **<0.001** | - | - |
| *Population-RM* |  | -8.9838 | -33.99603021 | 16.028446 | 0.475 | - | - |
| *Days since the 29th day of Drought x Population-RM* |  | 0.5308 | 0.05469984 | 1.006928 | **0.030** | - | - |
| *Stem RWC = Days since the 29th day of Drought x Population* | LM |  |  |  | **<0.001** | 55 | 0.73 |
| Intercept |  | 118.3478 | 104.167577 | 132.528118 | **<0.001** | - | - |
| *Days since the 29th day of Drought* |  | -1.6074 | -1.866141 | -1.348607 | **<0.001** | - | - |
| *Population-RM* |  | 12.9269 | 4.857052 | 20.996774 | **0.002** | - | - |
| *Root RWC = Days since the 29th day of Drought x Population* | LM |  |  |  | **<0.001** | 54 | 0.51 |
| Intercept |  | 97.7960 | 85.046141 | 110.5459402 | **<0.001** | - | - |
| *Days since the 29th day of Drought* |  | -0.8574 | -1.09040 | -0.6244183 | **<0.001** | - | - |
| *Population-RM* |  | 9.7382 | 2.421658 | 17.0547688 | **0.010** | - | - |

| **Model and Factors** | **Model type** | **Estimate** | **95% C.I. Estimates** | | ***p-value*** | **d.f. (res.)** | **Adjusted R square** |
| --- | --- | --- | --- | --- | --- | --- | --- |
|  |  |  | **2.5%** | **97.5%** |  |  |  |
| *Canopy Conductance = Days since the 29th day of Drought x Population* | LM |  |  |  | 0.219 | 54 | 0.03 |
| Intercept |  | 1.710e-05 | 2.844115e-06 | 3.136323e-05 | **0.020** | - | - |
| *Days since the 29th day of Drought* |  | -2.493e-07 | -5.207803e-07 | 2.208813e-08 | 0.071 | - | - |
| *Population-RM* |  | -1.971e-05 | -3.987444e-05 | 4.576837e-07 | 0.055 | - | - |
| *Days since the 29th day of Drought x Population-RM* |  | 3.430e-07 | -4.088858e-08 | 7.268434e-07 | 0.079 | - | - |
| *Probability of Mortality/100 = Days since the 29th day of Drought x Population* | GLM |  |  |  | **<0.001** | 57 | NA |
| Intercept |  | -10.04974 | -16.279168 | -5.8607657 | **<0.001** | - | - |
| *Days since the 29th day of Drought* |  | 0.20847 | 0.123628 | 0.3383138 | **<0.001** | - | - |
| *Population-RM* |  | -2.76760 | -5.752685 | -0.7046645 | **0.024** | - | - |
