## Supplementary material for "Relative water content consistently predicts drought mortality risk in seedling populations with different morphology, physiology, and times to death": Table S4

**Table S4.** Linear models used to predict changes in RWC over time and residual models testing the effects of morphology and physiology on population differences in dehydration rates. Data used corresponds to values starting at day 29 since the onset of drought to the end of the drought. Population categories correspond to North Plateau (NP) and Rocky mountain (RM).

| **Model and Factors** | **Estimate** | **95% C.I. Estimates** | | ***p-value*** | **d.f. (res.)** | **Adjusted R square** | **AICc** |
| --- | --- | --- | --- | --- | --- | --- | --- |
|  |  | **2.5%** | **97.5%** |  |  |  |  |
| *Plant RWC = Days since Onset of Drought* |  |  |  | **<0.001** | 45 | 0.59 | - |
| Intercept | 112.2879 | 97.627177 | 126.9487100 | **<0.001** | - | - | - |
| *Days since Onset of Drought* | -1.1199 | -1.388956 | -0.8508492 | **<0.001** | - | - | - |
| *Residuals Plant RWC = Population x Wood PLC x Plant NSC x Stomatal Conductance x Respiration* |  |  |  | **<0.001** | 39 | 0.46 | 388.77 |
| Intercept | -14.97221 | -29.9665 | 0.02208 | **0.05** | - | - | - |
| *Population RM* | -2.13277 | -16.90265 | 12.63711 | 0.772 | - | - | - |
| *Wood PLC* | 0.23308 | -0.05961 | 0.52576 | 0.115 | - | - | - |
| *Stomatal Conductance* | -1548.7176 | -4824.22847 | 1726.79324 | 0.345 | - | - | - |
| *Respiration* | -18.30396 | -30.62511 | -5.9828 | **0.005** | - | - | - |
| *Population RM x Wood PLC* | -0.08523 | -0.45894 | 0.28849 | 0.647 | - | - | - |
| *Population RM x Stomatal Conductance* | 7328.03463 | 2803.35297 | 11852.71629 | **0.002** | - | - | - |
| *Wood PLC x Stomatal Conductance* | 50.45403 | -20.92627 | 121.83432 | 0.161 | - | - | - |
| *Wood PLC x Respiration* | 0.55021 | 0.22837 | 0.87205 | **0.001** | - | - | - |
| *Population RM x Wood PLC x Stomatal Conductance* | -259.28753 | -398.11432 | -120.46075 | **<0.001** | - | - | - |
| *Residuals Plant RWC = Population x log(Root to shoot ratio) x log(Plant biomass)* |  |  |  | **<0.001** | 46 | 0.32 | 388.94 |
| Intercept | 11.704 | 4.340938 | 19.066895 | **0.003** | - | - | - |
| *log(Plant biomass)* | -9.078 | -16.156395 | -1.999588 | **0.013** | - | - | - |
| *log(Root to shoot ratio)* | -13.090 | -20.922028 | -5.257927 | **0.002** | - | - | - |
