## Supplementary material for "Relative water content consistently predicts drought mortality risk in seedling populations with different morphology, physiology, and times to death": Table S5

**Table S5.** Logistic models assessing the ability to predict mortality of RWC and water potential in each organ within a population.

| **Model and Factors** | | **Estimate** | **95% C.I. Estimates** | | ***p-value*** | **d.f. (res.)** | **V.E.** | **AIC** |
| --- | --- | --- | --- | --- | --- | --- | --- | --- |
|  |  |  | **2.5%** | **97.5%** |  |  |  |  |
|  | ***North Plateau*** | | | | | | | |
| *Probability of Mortality/100 = Root RWC* | |  |  |  | **0.001** | 34 | 0.54 | 31.80 |
| Intercept | | 4.98366 | 2.2091939 | 8.678722 | **0.002** | - | - | - |
| *Root RWC* | | -0.09276 | -0.1639429 | -0.044972 | **0.001** | - | - | - |
| *Probability of Mortality/100 = Stem RWC* | |  |  |  | **< 0.001** | 50 | 0.95 | 8.47 |
| Intercept | | 4.26961 | 2.0814809 | 8.46725910 | **0.003** | - | - | - |
| *Stem RWC* | | -0.10784 | -0.1846087 | -0.06349933 | **< 0.001** | - | - | - |
| *Probability of Mortality/100 = Leaf RWC* | |  |  |  | **< 0.001** | 48 | 0.95 | 8.88 |
| Intercept | | 4.95464 | 2.4975909 | 9.43711623 | **0.002** | - | - | - |
| *Leaf RWC* | | -0.10333 | -0.1772533 | -0.06096504 | **< 0.001** | - | - | - |
| *Probability of Mortality/100 = Plant RWC* | |  |  |  | **0.002** | 39 | 0.87 | 15.42 |
| Intercept | | 6.47419 | 3.2503647 | 12.0944653 | **0.002** | - | - | - |
| *Plant RWC* | | -0.13098 | -0.2491567 | -0.0686114 | **0.002** | - | - | - |
|  | ***Rocky Mountain*** | | | | | | | |
| *Probability of Mortality/100 = Root RWC* | |  |  |  | **0.005** | 35 | 0.54 | 28.08 |
| Intercept | | 4.78117 | 1.3645408 | 9.33191421 | **0.015** | - | - | - |
| *Root RWC* | | -0.09411 | -0.1745279 | -0.03855422 | **0.005** | - | - | - |
| *Probability of Mortality/100 = Stem RWC* | |  |  |  | **< 0.001** | 49 | 0.71 | 26.29 |
| Intercept | | 1.51439 | 0.09518901 | 3.15167840 | **0.046** | - | - | - |
| *Stem RWC* | | -0.06163 | -0.10373380 | -0.03298452 | **< 0.001** | - | - | - |
| *Probability of Mortality/100 = Leaf RWC* | |  |  |  | **< 0.001** | 50 | 0.61 | 29.67 |
| Intercept | | 2.74463 | 0.6939106 | 5.2292095 | **0.015** | - | - | - |
| *Leaf RWC* | | -0.06500 | -0.1064305 | -0.0337169 | **< 0.001** | - | - | - |
| *Probability of Mortality/100 = Plant RWC* | |  |  |  | **< 0.001** | 43 | 0.67 | 26.43 |
| Intercept | | 3.84161 | 1.2997532 | 7.12454758 | **0.008** | - | - | - |
| *Plant RWC* | | -0.08141 | -0.1396342 | -0.04074093 | **< 0.001** | - | - | - |
| **Model and Factors** | | **Estimate** | **95% C.I. Estimates** | | ***p-value*** | **d.f. (res.)** | **V.E.** | **AIC** |
|  |  |  | **2.5%** | **97.5%** |  |  |  |  |
|  | ***North Plateau*** | | | | | | | |
| *Probability of Mortality/100 = Soil Water Potential* | |  |  |  | **<0.001** | 53 | 0.59 | 35.66 |
| Intercept | | -3.9644 | -6.842051 | -2.2048206 | **<0.001** | - | - | - |
| *Soil Water Potential* | | -1.3502 | -2.366627 | -0.7167899 | **<0.001** | - | - | - |
| *Probability of Mortality/100 = Stem Water Potential* | |  |  |  | **<0.001** | 53 | 0.93 | 11.46 |
| Intercept | | -6.5164 | -12.347141 | -3.5992918 | **0.002** | - | - | - |
| *Stem Water Potential* | | -1.2105 | -2.188772 | -0.6825154 | **<0.001** | - | - | - |
| *Probability of Mortality/100 = Leaf Water Potential* | |  |  |  | **<0.001** | 52 | 0.89 | 14.34 |
| Intercept | | -6.758 | -13.420706 | -3.6954542 | **0.002** | - | - | - |
| *Leaf Water Potential* | | -1.195 | -2.227083 | -0.6702828 | **<0.001** | - | - | - |
|  | ***Rocky Mountain*** | | | | | | | |
| *Probability of Mortality/100 = Soil Water Potential* | |  |  |  | **0.010** | 53 | 0.46 | 37.00 |
| Intercept | | -3.8623 | -6.974155 | -2.1278054 | **<0.001** | - | - | - |
| *Soil Water Potential* | | -1.1648 | -2.345810 | -0.4730897 | **0.010** | - | - | - |
| *Probability of Mortality/100 = Stem Water Potential* | |  |  |  | **0.002** | 52 | 0.88 | 19.37 |
| Intercept | | -5.7081 | -11.295333 | -3.193501 | **0.002** | - | - | - |
| *Stem Water Potential* | | -0.8257 | -1.624107 | -0.435619 | **0.002** | - | - | - |
| *Probability of Mortality/100 = Leaf Water Potential* | |  |  |  | **<0.001** | 52 | 0.80 | 22.78 |
| Intercept | | -5.1561 | -9.064376 | -3.0238930 | **<0.001** | - | - | - |
| *Leaf Water Potential* | | -0.7508 | -1.325623 | -0.4035579 | **<0.001** | - | - | - |
