## Supplementary material for "Relative water content consistently predicts drought mortality risk in seedling populations with different morphology, physiology, and times to death": Table S6

**Table S6.** Logistic models assessing the consistency of the relationships between probability of Mortality and relative water content (RWC) or water potential among populations and organs. Population categories correspond to North Plateau (NP) and Rocky mountain (RM). Organ categories correspond to leaf, stem, roots, whole-plant, and soil. We entered whole-plant and soil as an organ to be able to compare them to the rest of organs.

| **Model and Factors** | **Estimate** | **95% C.I. Estimates** | | ***p-value*** | **d.f. (res.)** | **AIC** |
| --- | --- | --- | --- | --- | --- | --- |
|  |  | **2.5%** | **97.5%** |  |  |  |
| *Probability of Mortality = RWC * Population * Organ* |  |  |  |  | 348 | 175.1 |
| Intercept | 4.954645 | 2.49759289 | 9.43711507 | **0.002** | - | - |
| *RWC* | -0.103329 | -0.17725331 | -0.06096500 | **< 0.001** | - | - |
| *Population-RM* | -2.210012 | -7.06192957 | 1.34636292 | 0.262 | - | - |
| *Organ-Plant* | 1.519549 | -3.97511538 | 7.68110342 | 0.566 | - | - |
| *Organ-Roots* | 0.029017 | -5.15107061 | 4.53011361 | 0.990 | - | - |
| *Organ-Stem* | -0.685038 | -5.68687207 | 4.16151679 | 0.753 | - | - |
| *RWC * Population-RM* | 0.038328 | -0.02155419 | 0.11789299 | 0.243 | - | - |
| *RWC * Organ-Plant* | -0.027653 | -0.15367542 | 0.06955652 | 0.588 | - | - |
| *RWC * Organ-Roots* | 0.010569 | -0.07288420 | 0.09767735 | 0.791 | - | - |
| *RWC * Organ-Stem* | -0.004513 | -0.09221156 | 0.08169024 | 0.910 | - | - |
| *Population-RM * Organ-Plant* | -0.422577 | -7.37617276 | 6.07617875 | 0.895 | - | - |
| *Population-RM * Organ-Roots* | 2.007525 | -4.13144001 | 8.82999003 | 0.531 | - | - |
| *Population-RM * Organ-Stem* | -0.545209 | -5.98431389 | 4.99210991 | 0.831 | - | - |
| *RWC * Population-RM * Organ-Plant* | 0.011241 | -0.10293724 | 0.14764083 | 0.849 | - | - |
| *RWC * Population-RM * Organ-Roots* | -0.039680 | -0.15626517 | 0.06767046 | 0.471 | - | - |
| *RWC * Population-RM * Organ-Stem* | 0.007881 | -0.08965499 | 0.10636449 | 0.866 | - | - |

| **Model and Factors** | **Estimate** | **95% C.I. Estimates** | | ***p-value*** | **d.f. (res.)** | **AIC** |
| --- | --- | --- | --- | --- | --- | --- |
|  |  | **2.5%** | **97.5%** |  |  |  |
| *Probability of Mortality = Water potential * Population * Organ* |  |  |  |  | 315 | 140.62 |
| Intercept | -6.75760 | -13.4207055 | -3.6954542 | **0.002** | - | - |
| *Water potential* | -1.19521 | -2.2270832 | -0.6702828 | **< 0.001** | - | - |
| *Population-RM* | 1.60149 | -3.4211186 | 8.6136058 | 0.543 | - | - |
| *Organ-Soil* | 2.79325 | -1.4895129 | 9.6921907 | 0.263 | - | - |
| *Organ-Stem* | 0.24124 | -6.4449649 | 7.5774654 | 0.937 | - | - |
| *Water potential * Population-RM* | 0.44437 | -0.3366592 | 1.5321003 | 0.291 | - | - |
| *Water potential * Organ-Soil* | -0.15503 | -1.3158572 | 1.0504650 | 0.775 | - | - |
| *Water potential * Organ-Stem* | -0.01526 | -1.1356876 | 1.1495089 | 0.976 | - | - |
| *Population-RM * Organ-Soil* | -1.49940 | -9.0635572 | 4.4072637 | 0.628 | - | - |
| *Population-RM * Organ-Stem* | -0.79325 | -9.4144049 | 7.0472438 | 0.835 | - | - |
| *Water Potential * Population-RM * Organ-Soil* | -0.25894 | -1.8746480 | 1.1901370 | 0.726 | - | - |
| *Water Potential * Population-RM * Organ-Stem* | -0.05958 | -1.4018790 | 1.2175489 | 0.923 | - | - |
